# Resource competition drives an invasion-replacement event among shrew species on an island

**DOI:** 10.1101/2022.04.27.489660

**Authors:** Samuel S. Browett, Rebecca Synnott, Denise B. O’Meara, Rachael E. Antwis, Stephen S. Browett, Kevin J. Bown, Owen S. Wangensteen, Deborah A. Dawson, Jeremy B. Searle, Jon M. Yearsley, Allan D. McDevitt

## Abstract

1. Invasive mammals are responsible for the majority of native species extinctions on islands. While most of these extinction events will be due to novel interactions between species (e.g. exotic predators and naive prey), it is more unusual to find incidences where a newly invasive species causes the decline/extinction of a native species on an island when they normally coexist elsewhere in their overlapping mainland ranges.
2. We investigated if resource competition between two insectivorous small mammals was playing a significant role in the rapid replacement of the native pygmy shrew (*Sorex minutus*) in the presence of the recently invading greater white-toothed shrew (*Crocidura russula*) on the island of Ireland.
3. We used DNA metabarcoding of gut contents from >300 individuals of both species to determine each species’ diet and measured the size of individuals (weight and length) during different stages of the invasion in Ireland (before, during and after the species come into contact with one another) and on a French island where both species have long coexisted (acting as a natural ‘control’ site). Dietary composition, niche width and overlap and size were compared in these different stages.
4. The size of the invasive *C. russula* and composition of its diet changes between when it first invades an area and after it becomes established. Individuals are larger and they consume larger invertebrates at the invasion front, before switching towards the smaller prey taxa that are more essential for the survival of the native species after establishment. As a result, the level of interspecific dietary overlap increases from between 11–14% when they first come into contact with each other to between 39–46% after the invasion.
5. Here we show that an invasive species can quickly alter its dietary niche in a new environment, leading to negative impacts that were not previously predicted based on the coexistence of these species in other parts of their mainland ranges. As well as causing the replacement of a native small mammal, the invasive shrew may be rapidly exhausting local resources of larger invertebrate species. These subsequent changes in terrestrial invertebrate communities could have severe impacts further downstream on ecosystem functioning and services.

## Introduction

The rate at which species are introduced into novel, non-native ranges has been accelerating due to increased globalisation (Hulme, 2009; Seebens et al., 2017). As well as increasing societal impacts associated with the management costs of these invasions (Diagne *et al*., 2021), the ecological impacts that newly introduced invasive species can have on local ecosystems are now so severe that they are considered the most common cause of vertebrate extinctions (Bellard *et al*., 2016a). This is particularly evident in island ecosystems, where invasive species are responsible for most of the damage/impacts caused to local fauna and flora (Bellard *et al*., 2016a, b; Spatz *et al*., 2017). Islands often have more simplified ecological systems with smaller species communities than their mainland counterparts and therefore tend to be more susceptible to anthropogenic impacts (Spatz *et al*., 2017). Invasive species have been implicated in >80% of species extinctions on islands over the last 500 years (Bellard *et al*., 2016a), and invasive mammals are responsible for the majority of these (Jones *et al*. 2016).

The impacts of species invasions can either be through direct (e.g. predation Doherty *et al*., 2016) or indirect (e.g. competition or trophic cascades; Hernandez-Brito *et al*., 2018; Benkwitt *et al*., 2021) mechanisms. The strength of any competitive interaction between invasive and native species may depend on the community composition and environment of the invaded area, the speed of the invasion, and potential trade-offs between dispersal, reproduction and competitive ability of the invasive species as it expands its range (Burton *et al*., 2010). However, predicting the impacts of non-native species on novel ecosystems can be challenging (Griffen *et al*., 2020), particularly when a newly introduced non-native species coexists with components of the novel community elsewhere in its range (McDevitt *et al*., 2014).

The recent introduction of the greater white-toothed shrew (*Crocidura russula*) into the island of Ireland provides an example of how unpredictable the impacts of species invasions can be on the local fauna. *Crocidura russula* was likely accidentally introduced into Ireland in the early 2000s via horticultural imports from mainland France (Tosh *et al*., 2008; McDevitt *et al*. 2014; O’Meara *et al*., 2014; Gargan *et al*., 2016). Prior to its arrival, the pygmy shrew (*Sorex minutus*) was the only species of shrew present in Ireland. *Sorex minutus* is sympatric with multiple shrew species across the European mainland, including *C*. *russula*. Differential resource use and niche separation among these insectivorous small mammals is known to be integral for facilitating multi-shrew communities (Rey, Noguerales and García-Navas, 2019) and this has been proposed to facilitate the sympatric existence of *S. minutus* (albeit in low abundance) with larger species of shrews in mainland Europe (Churchfield and Rychlik, 2006). Indeed, *S. minutus* and *C. russula* are the only shrew species present on the small island of Belle Île off northwestern France and coexist with one another in large numbers amongst a similar small mammal community that is present in Ireland (McDevitt *et al*., 2014, Gargan *et al*., 2016). In contrast to this however, the invasion and rapid spread of *C. russula* in Ireland is clearly associated with the local disappearance of *S. minutus* (Montgomery, Lundy and Reid, 2012; McDevitt *et al*., 2014; Montgomery, Montgomery and Reid, 2015). Although *C. russula* is known to harbour a novel strain of pathogenic *Leptospira* and while the potential role of novel pathogens/disease in this invasion-replacement event cannot be completely discounted, no evidence of disease onset was apparent after experimental infections (Nally *et al*., 2016). *Crocidura russula* is known to outcompete other shrew species when it colonises an area/island in other regions (Biedma *et al*., 2018; Cornette *et al*. 2015) but the exact mechanism(s) of how this occurs is uncertain. Therefore, McDevitt *et al*. (2014) proposed that *S. minutus* may have experienced a competitive release on the island in the absence of other shrew species and is now not able to adapt quickly enough to a new invasive competitor.

This recent and ongoing invasion therefore presents us with a unique opportunity to examine resource competition between a native species and an invasive competitor before, during, and after an invasion in a real-time setting. There is a narrow region at the edge of the *C. russula* invasive range in Ireland where both shrew species overlap temporarily until *S. minutus* disappears in as little as a year (**Fig.1**; McDevitt *et al*., 2014). Further inside the well-established invasive range of *C. russula*, there is no evidence that *S. minutus* is still present (Montgomery *et al*., 2012, 2015; McDevitt *et al*., 2014; this study). The goal of this study was to investigate the diet and size (determined by length and weight) of both species of shrews at different stages of the invasion (**Fig. 1**). In addition, the diet and size of both species was investigated in Belle Île, where both species co-exist. Belle Île is an ideal natural ‘control’ site as the habitat types are similar to Ireland and it has the same small mammal community present (*C. russula*, *S. minutus*, *Clethrionomys glareolus* (bank vole) and *Apodemus sylvaticus* (wood mouse); McDevitt *et al*., 2014). To determine if resource competition is a contributing factor in the local replacement of *S. minutus* in response to the *C. russula* invasion in Ireland, DNA metabarcoding (Pompanon *et al*., 2012; Deagle *et a*l., 2019) was applied to the gut contents of shrews to characterise the diet of *S. minutus* and *C. russula* at different stages of the invasion in Ireland and Belle Île in order to investigate the levels of dietary overlap and interspecific competition between them.

**Figure 1.**
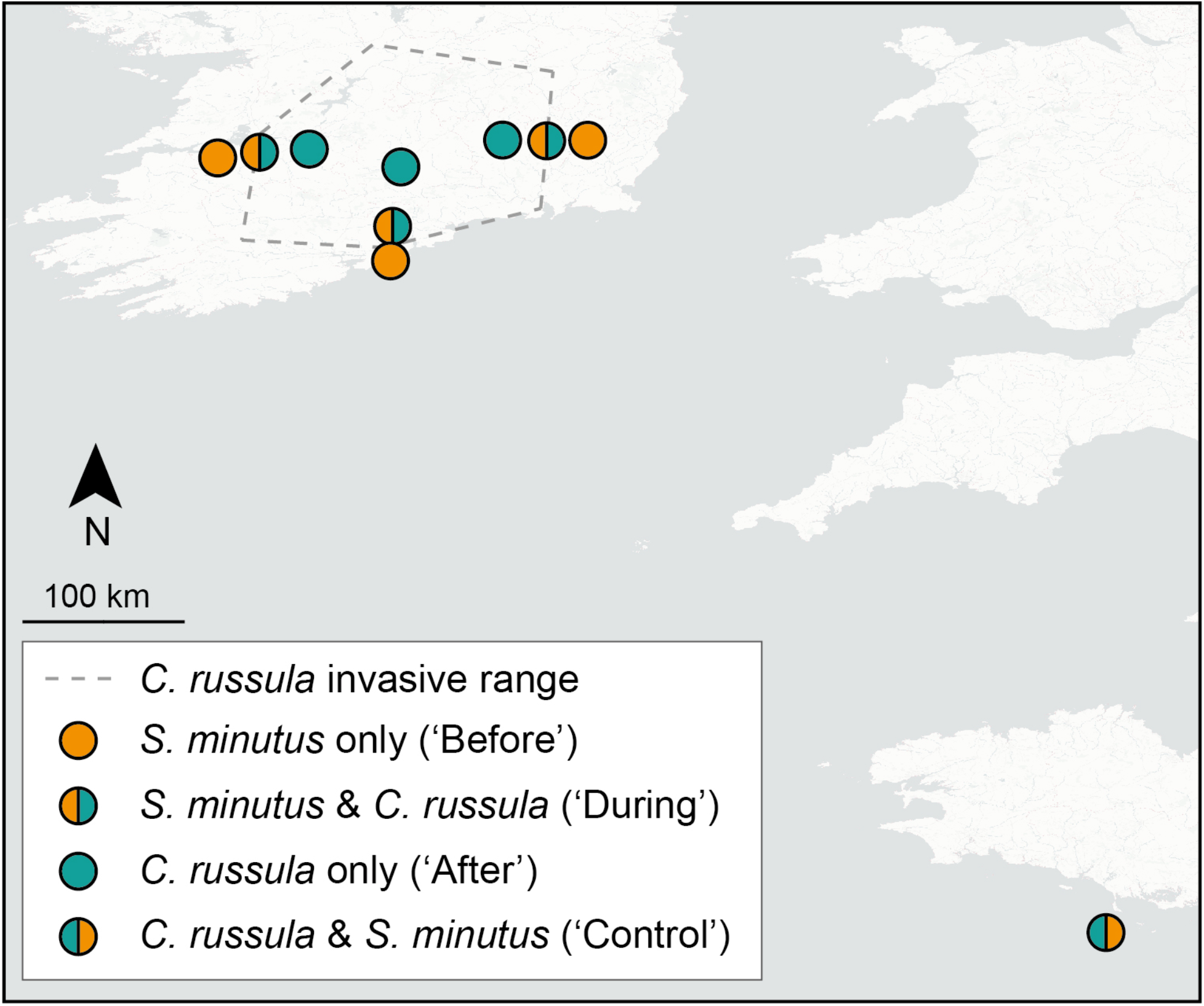
Sampling sites in each of three transects covering the different invasion stages (at the time of sampling) of *Crocidura russula* in Ireland (‘Before’, ‘During’ and ‘After’) and the ‘Control’ site in Belle Île. The presence of *C. russula* is indicated in green and *Sorex minutus* in orange.

## Methods

### Study design

Sampling sites were chosen to investigate the impacts of the spread of *C. russula* in Ireland. The south of Ireland was divided into three ‘stages’ of the invasion: ‘before’, ‘during’ and ‘after’ from the perspective of the impacted native *S. minutus* (**Fig. 1**). ‘Before’ the invasion denotes where *C. russula* has not yet invaded and with only *S. minutus* present. ‘During’ the invasion is at the edge of the invasive range where *C. russula* first comes into contact with *S. minutus* and therefore both species overlap. ‘After’ the invasion is where only *C. russula* is present and *S. minutus* is no longer observed. To accommodate for geographical variation of available prey, shrews were sampled along eastern, western and southern transects through the invasive range (**Fig. 1**). Due to the small size of the ‘control’ site in Belle Île (84 km^2^) where both species overlap, shrews were sampled from across the entire island.

To account for seasonal variation in diet (Grainger and Fairley, 1978), sampling was conducted over two seasonal time periods. The first seasonal sampling period took place from 19/08/2017 to 17/10/2017, referred to hereafter as ‘summer’. The second sampling period took place from 16/02/2018 to 06/04/2018, referred to hereafter as ‘spring’. These dates by-pass peak breeding months and should target the same cohort of shrews across the year.

### Sample Collection

Trap sites were chosen at accessible hedgerows along secondary and tertiary roads adjacent to agricultural land (pasture or arable). Shrews were trapped using trip-traps (Proctor Bros. Ltd., UK) with no bait (to avoid interference with dietary analyses). See Supplementary Material and **Fig. S1** for more detail on trapping and sites. Shrews were immediately euthanised by cervical dislocation following guidelines set out by Sikes (2016). All trapping and procedures were performed under the appropriate licences C21/2017 (National Parks and Wildlife Service, Ireland), AE18982/I323 (Health Products Regulatory Authority; Ireland) and A-75-1977 (Belle Île, France), and ethical approvals ST1617-55 (University of Salford, UK) and AREC-17-14 (University College Dublin, Ireland). Male and female adults were sampled. Each shrew was weighed using a 50g Pesola spring scale and body length was measured for each sample. The gut tract (stomach and intestines) was removed and stored in absolute ethanol at a 1:4 (sample:ethanol) ratio (Egeter, Bishop and Robertson, 2015). To avoid cross-contamination, all dissections were performed on disposable bench covers and all tools were cleaned and flamed between samples. Gut contents were stored in ethanol at -20°C upon returning from the field to the lab (max. 12 days). A total of 99 *S. minutus* and 124 *C. russula* were caught from Ireland and a total of 40 *C. russula* and 40 *S. minutus* were caught from Belle Île (see **Table S1** for sample sizes by transect and zone).

### Lab Protocols

DNA was extracted from the gut contents using the Qiagen PowerSoil Kit (Qiagen Ltd.), with five extraction blanks. A 133bp fragment of the mtDNA COI gene was amplified from DNA extracts using the primers LepF1 (5′-ATTCHACDAAYCAYAARGAYATYGG-3′) and EPT-long-univR (5′-ACTATAAAARAAAATYTDAYAAADGCRTG-3′; Gillet *et al*., 2015)) according to the protocol described in Browett *et al*. (2021). These primers were previously shown to amplify a wide range of prey taxa in shrews (Browett *et al.,* 2021). The final library with a total of 303 samples, five extraction blanks and 20 PCR blanks was sequenced on two Illumina MiSeq runs using V2 2x150 bp cycle kits, both loaded at 9pM with a 5% PhiX (v3, Illumina) spike. See the Supplementary Material for more detailed DNA extractions, PCR, library preparation and sequencing information.

### Bioinformatics and data filtering

Processing of raw sequence reads was performed using the Obitools metabarcoding software pipeline (Boyer *et al*., 2016). After aligning the paired-end reads, sequences with an alignment quality score >40 and a length between 128–138bp were retained (Browett *et al*., 2021). See the Supplementary Material for further details of the bioinformatics undertaken. All MOTUs belonging to non-prey taxa (e.g. vertebrates and parasites) were removed. Samples with less than 1000 prey reads were removed. To mitigate false positive detections, MOTUs were removed from each sample if they were represented by less than 0.1% of the total prey reads in that individual sample (Alberdi *et al*., 2018; Deagle *et al*., 2019).

To determine the coverage of samples, rarefaction curves and species accumulation curves were generated using the R package vegan (Oksanen *et al*., 2019). In addition, the *depth_cov()* function in the *hilldiv* R package (Alberdi and Gilbert, 2019) was used to clarify if sufficient read depth was obtained for each sample, using the qvalue = 1 (equivalent to Shannon diversity measure). A second dataset containing a ‘core’ diet was created by removing rare prey taxa found in a single sample. This strategy is recommended for dietary studies, particularly for calculating resource overlap values (Brown *et al*., 2014; Arrizabalaga-Escudero *et al*., 2018).

### Diet Composition and Niche Overlap and Width

Three methods for quantifying the importance of different taxa to a population’s diet were compared as described and recommended by Deagle *et al*. (2019). These were relative read abundance (RRA), percentage of occurrence (POO) and weighted percentage of occurence (wPOO). To determine the compositional difference in prey taxa identified between different invasion stages in Ireland and Belle Ile, PERMANOVA’s were performed using the *adonis()* function in the vegan package in R (Oksanen *et al*., 2019). The multivariate distances of samples to the group centroid were calculated using *betadisper()* function in the vegan package and a permutation test for homogeneity of multivariate dispersions was used to determine if there was a similar level of variance between each group. These distances were calculated using reads transformed into relative read abundances and using the Bray-Curtis distance metric (using relative read abundance), and the Jaccard distance metric (using presence-absence). These were performed for prey taxa grouped at MOTU, species, genus, family and order levels. NMDS plots were generated to visualise beta diversity measures, with enough dimensions to reduce stress to approximately 0.1.

The Pianka’s (1973) niche overlap index (Ojk) was calculated in the R package *ecosimR* (Gotelli and Ellison, 2013) to identify overlap in diet between *S. minutus* and *C. russula* in different study sites. To determine if resource overlap was significantly higher or lower than expected, a null model was created by running 10,000 resource utilization simulations using randomisation function RA3 which reshuffles values within each predator group. Observed overlap values were compared to this null model to determine if the observed overlap is more or less than a random situation. The samples were grouped according to shrew species, Ireland vs Belle Île, and invasion stage. The niche width of each group was measured using the standardised Levin’s index and Shannon diversity measure (for details on measurements see Razgour *et al*., 2011) on the POO values for each group using the R package *spaa* (Zhang, 2016). If there was a large difference in sample size between groups, larger sample sizes were randomly subsampled to the same as the smallest group 50 times and the average diversity scores were recorded.

## Results

### Body Measurements

The Belle Île population of *S. minutus* had larger individuals (mean ± SE weight: 4.36 ± 0.12 g; length: 96.47 ± 0.84 mm) compared to individuals of the same species sampled in Ireland (weight: 3.34 ± 0.05 g; length 91.42 ± 0.45 mm; ANOVA post hoc Tukey p < 0.0001 for both weight and length; **Table S2**; **Figs. 2B** **and S2**). There is no evidence of a size difference between Irish *S. minutus* sampled ‘before’ the invasion (weight: 3.28 ± 0.06 g; length 90.62 ± 0.61 mm) and ‘during’ it (weight: 3.40 ± 0.07 g; length: 92.28 ± 0.66 mm). Conversely, *C. russula* in Ireland are larger compared to those in Belle Île. The *C. russula* sampled ‘during’ the invasion in Ireland were the largest sampled (weight: 11.45 ± 0.25 g; length: 117.73 ± 0.56 mm) when compared to shrews sampled ‘after’ the invasion (weight: 10.58 ± 0.17 g; length: 114.73 ± 0.44 mm; p < 0.0001) and when compared to Belle Île *C. russula* (weight: 9.84 ± 0.2 g; length: 111.81 ± 0.78 mm; p < 0.002; **Table S2**; **Figs. 2B** **and S2**). These overall patterns in weight and length were generally evident within a sampling season also but were more pronounced in samples collected in the spring period (**Table S2**; **Fig. S3**).

**Figure 2.**
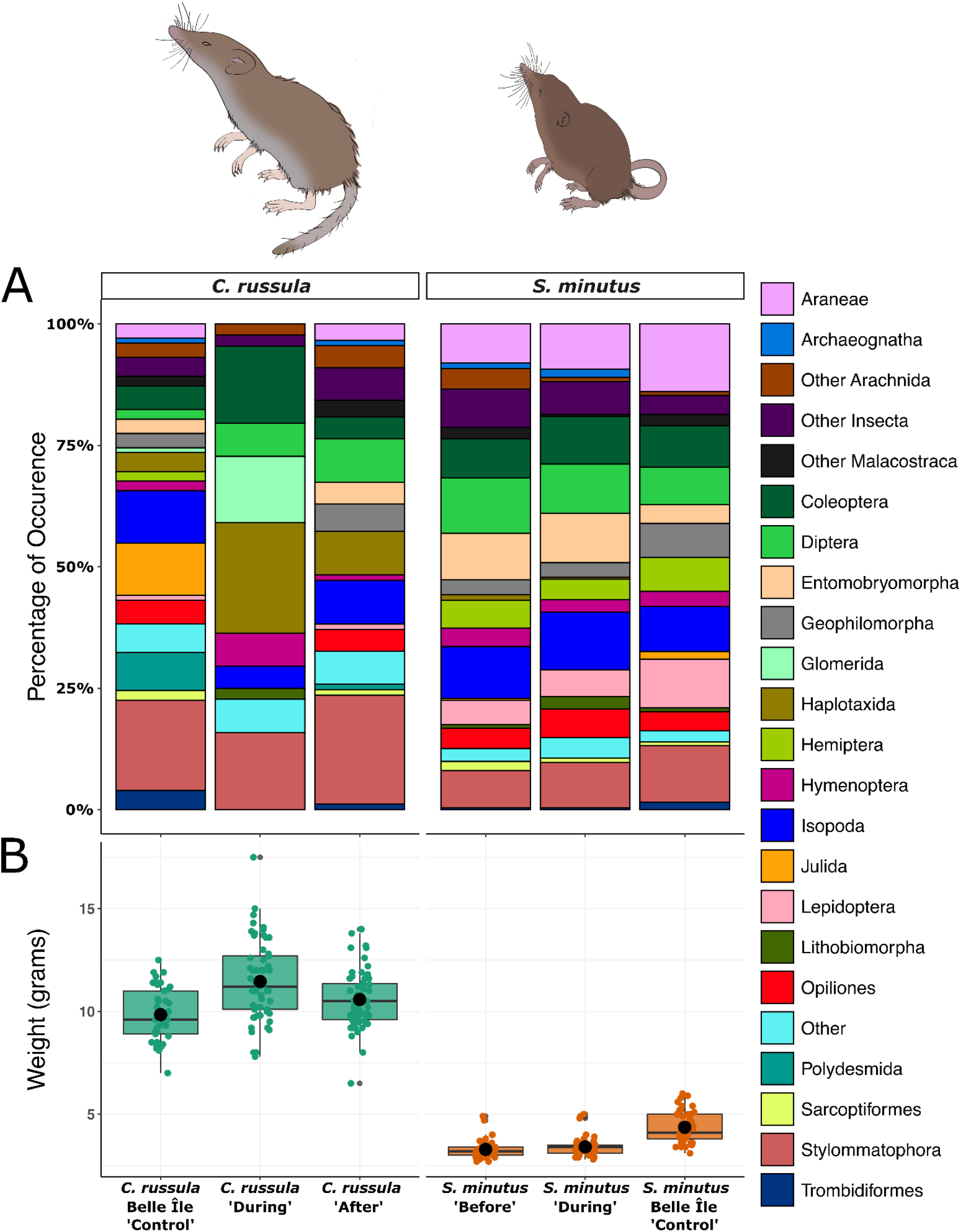
The composition of *Crocidura russula* and *Sorex minutus* diet (using Percentage of Occurrence or POO) grouped to the level of invertebrate order (each represented by a unique colour) and for each invasion stage in Ireland (‘Before’, ‘During’ and ‘After’) and the ‘Control’ site in Belle Île (A). The weights of *C. russula* (green) and *S. minutus* (orange) for each invasion stage in Ireland and Belle Île (B).

### Sequencing

The two sequencing runs generated a total of 30,172,418 reads. After quality filtering of sequences and chimera removal, there were 21,091,503 reads across the 303 samples (average read depth of 68,459 reads per sample) and 25 negative controls. A full breakdown of retained sequences is provided in **Table S3**. The dataset utilising the sequence clustering threshold at 98% similarity yielded 33,801 non-singleton MOTUs. There were a total of 38,535 reads (0.18% of total reads) from 394 MOTUs identified in the negative controls. The collective read count of MOTUs in the negative controls ranged from 1 to 14,606. The most prominent reads in the negative controls were from the family Soricidae (shrews) (**Fig. S4**), most likely due to a combination of strong host amplification and ‘tag jumping’ (Schnell *et al*., 2015). Host amplification ranged between ∼15.6% and ∼99.95% in *C. russula* and between 0.14% and ∼99% in *S. minutus*.

The final dataset of prey items contained 994 MOTUs across 178 samples (59 *C. russula* and 119 *S. minutus*) with an average read depth of 34,955 prey reads per individual. Multiple samples of both species had notably less food remains inside the GI tract during dissection, which subsequently only returned host reads and the samples were filtered out. Previous work on *C. russula* showed that using alternative primer sets that do not amplify vertebrate DNA will still result in sample drop outs due to empty GI tracts (Browett *et al.,* 2021). The sequencing depth showed sufficient coverage for each of the 178 samples, with richness (q=0) showing 100% coverage and eveness (q=1) values of >98%. Species accumulation curves show that at the species/MOTU level, the plateau was not reached for either species in either Ireland or Belle Île (see **Fig. S5**). Sample coverage improves when agglomerating taxa to higher levels, with a plateau reached at the order level. This is a common feature of dietary metabarcoding studies in insectivorous species (Tournayre *et al*., 2020).

### Diet Composition

A wide range of invertebrate taxa are consumed by both species (**Figs. S6 and S7**). When using RRA at the level of the taxonomic order of the prey for each individual shrew (**Fig. S8**), it is evident that there is wide variation between individuals. This observed variation is consistent with the high level of variation between samples seen in the beta diversity results. RRA, POO and wPOO performed very similarly for quantifying the diet of groups of shrew samples (**Fig. S9**), with discrepancies between certain prey orders (see Supplementary Material for more details). The amplification biases seen between certain primers and prey taxonomic groups (Krehenwinkel *et al*., 2017; Bista *et al*., 2018) means that using the wPOO or POO metrics are the more conservative approach. There was little difference between these two metrics, and therefore only the POO values are reported in the main text while the RRA metrics are available in the Supplementary Material for individuals. POO reveals a similar diet composition between Irish and Belle Île *S. minutus*. The Irish *S. minutus* population has a higher proportion of Diptera, Enterobryomorpha and Isopoda, while Belle Île *S. minutus*l shows a higher rate of predation on Araneae (**Fig. 2A**). Visually the composition of the *C. russula* diet differs for prey orders between Belle Île and the two invasion stages within Ireland, complementing the PERMANOVA results (**Table 1**). The composition of *C. russula* ‘after’ the invasion has the closest resemblance to the *S. minutus* diet in Ireland.

**Table 1.**
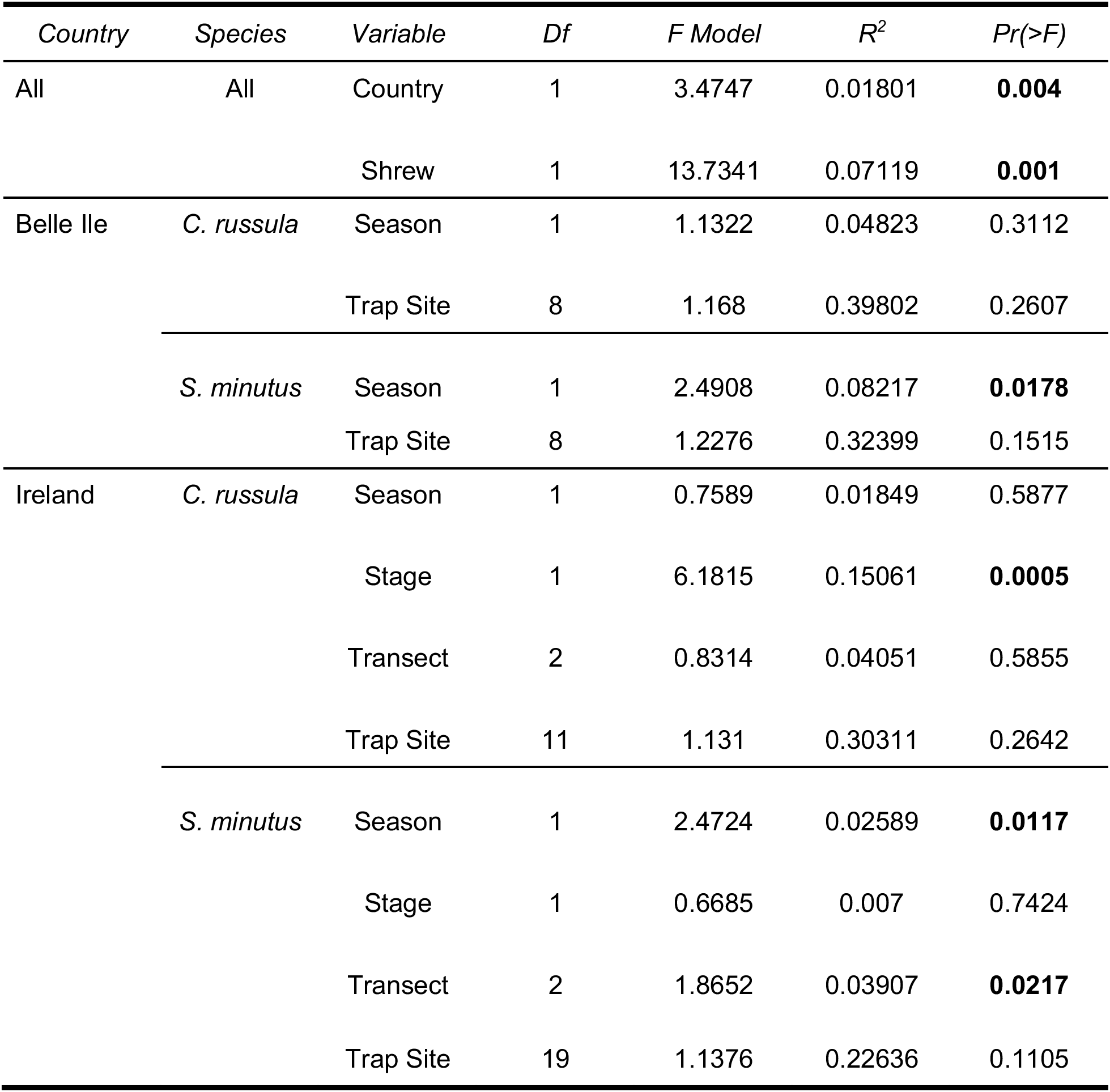
PERMANOVA results at the order level of identified prey taxa to show the prey composition dissimilarities using the Bray-Curtis distance method. Shrews are grouped according to species and island of capture. A PERMANOVA is performed on each group, examining multiple variables (Season, Trap Site, Invasion Stage and Transect) that contribute to compositional variance within each group. Variables are treated sequentially from top to bottom in each group. Significant values are indicated in bold (< 0.05).

Many prey orders appear to remain consistent throughout the year, such as slugs and snails (Stylommatophora) and earthworms (Haplotaxida) in *C. russula* in Ireland (**Fig. S10**). *Sorex minutus* shows the most notable seasonal shifts in prey orders (also shown by the PERMANOVA results; **Table 1**). *S. minutus* in Belle Île shows a decrease in consumption of Hemiptera in the spring with a dramatic increase in consumption of Araneae and Lepidoptera during the summer sampling period (**Fig. S10**).

### Niche Width and Overlap

The standardised Levin’s index indicates that the niche width of both *S. minutus* and *C. russula* are similar (**Table S4**). The standardised Levin’s index indicates that the niche width of Irish *C. russula* is wider than Belle Île *C. russula* at the MOTU (0.59 and 0.48), species (0.55 and 0.44) and genus (0.47 and 0.42 respectively) level but with similar niche width at the family level (0.44 and 0.43). However, the niche width at the order level is narrower for Irish *C. russula* compared to Belle Île (0.53 and 0.60 respectively). The Irish *S. minutus* population show a narrower niche width compared to Belle Île at the MOTU (0.50 and 0.58), species (0.45 and 0.54), genus (0.43 and 0.55) and family (0.48 and 0.59) level but a similar niche width at the order level (0.60 and 0.58 respectively; **Table S4**).

The PERMANOVA shows there is a difference in the composition of the diet at the MOTU level between shrew species (R^2^ = 0.02, p = 0.001), and between Ireland and Belle Île (R^2^ = 0.02, p = 0.001). Among the top 20 MOTUs contributing most to the differences between shrews and between Ireland and Belle Île, there are mostly MOTUs belonging to Gastropoda (slugs and snails), Clitellata (worms) and Diplopoda (millipedes). There are notable differences in proportions of these orders in the diet of both shrews (**Fig. 2A**). While PERMANOVAs showed no difference between Irish *C. russula* according to the season, transect or trap site, there was a small difference according to the invasion stage (R^2^ = 0.05, p = 0.029). This difference is primarily caused by MOTUs from Insecta and Gastropoda. Season also showed no effect in Belle Île *C. russula*, but there was an observed difference between trap sites within the island (R^2^ = 0.45, p = 0.001).

Irish *S. minutus* show a change in dietary composition according to season (R2 = 0.02, p = 0.001), transect (R2 = 0.04, p = 0.001) and trap site (R2 = 0.23, p = 0.001). The majority of MOTUs contributing to the differences in transect and sampling sites are the same, primarily belonging to Coleoptera and Lepidoptera. The difference occurring between seasons is primarily driven by MOTUs ascribed to the Insecta class. Belle Île *S. minutus* also show slight differences in the composition of their diet between seasons (R2 = 0.08, p = 0.001), but not trap sites. This shift in seasonal diet is primarily influenced by Insecta and Arachnida, which is noticeable in compositional change using POO measures (see **Fig. S10**).

The NMDS plot shows that while all four stages between two species may be different in their centroid/core diet, there is still considerable overlap between samples (**Fig. 3**). These patterns complement the PERMANOVA results, showing differences between groups but with low R^2^ values that indicate that the tested variables explain less than 10% of the variation, except trap sites explaining up to 32% of the variation in *S. minutus* and 40% in *C. russula*. When considering only the Irish samples at different invasion stages, there is still considerable overlap. However, as MOTUs are agglomerated in genus, family and order, the *C. russula* sampled ‘during’ the invasion appear the most different, confirming the PERMANOVA results demonstrating that invasion stage explains most of the variation in the diet of Irish *C. russula*. These plots suggest a higher similarity in diet between *C. russula* captured ‘after’ the invasion and *S. minutus* ‘before’ the invasion, particularly when prey species are grouped to higher taxonomic levels (**Fig. 3**).

**Figure 3.**
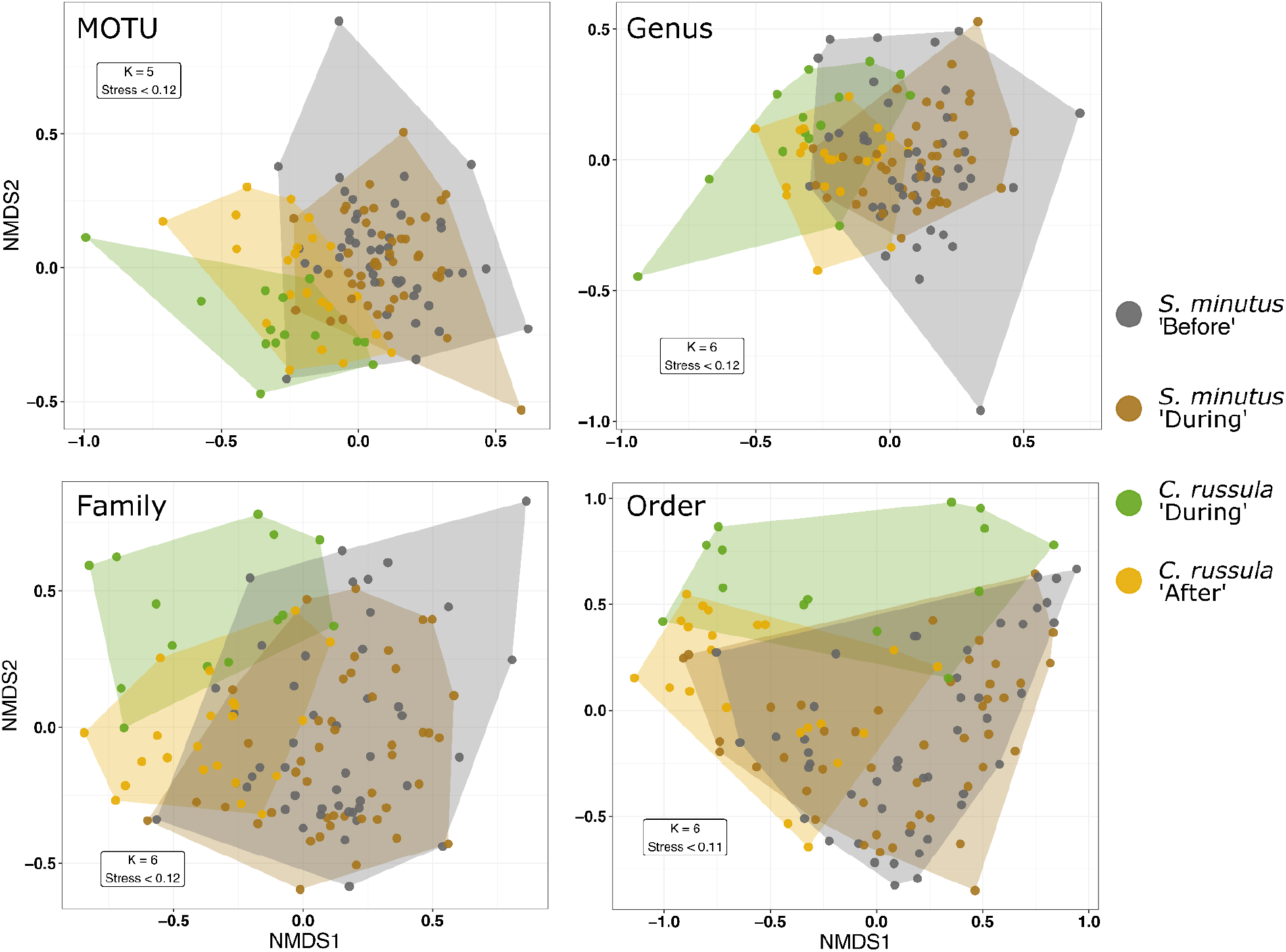
NMDS plot generated to visualise dietary overlap using the Bray-Curtis dissimilarity method using prey at the (clockwise from top left) Molecular Operational Taxonomic Unit (MOTU), genus, order and family level according to the invasion stages (‘Before’, ‘During’ and ‘After’; see main text) in Ireland for each species. Note there is less overlap between *C. russula* during the invasion (green) and the rest of the groups (corresponding to the dietary overlap results; Table 2)

**Table 2.** Dietary overlap (Pianka index) values using Percentage of Occurrence (POO) and including all Molecular Operational Taxonomic Units (All MOTUs) and Core MOTUs (removing rare prey taxa found in a single sample). This includes comparisons between islands (Ireland and Belle Île) and the ‘Before’, ‘During’ and ‘After’ stages of *Crocidura russula*’s invasion of Ireland (see main text). Index values range from 0 (no overlap) to 1 (complete overlap). Significance indicated at *0.05; **0.01.

| *Interspecific and Island Comparisons* |  |  | *All MOTUs* | *Core MOTUs* |
| --- | --- | --- | --- | --- |
| *C. russula* - Belle Île | vs | *C. russula* - Ireland | 0.42877* | 0.45655 |
| *C. russula* - Belle Île | vs | *S. minutus* - Belle Île | 0.4617* | 0.50289 |
| *C. russula* - Ireland | vs | *S. minutus* - Ireland | 0.35772** | 0.37765 |
| *S. minutus* - Belle Île | vs | *S. minutus* - Ireland | 0.39034** | 0.47217* |
| *C. russula* - ‘During’ | vs | *S. minutus* - ‘During’ | 0.11326 | 0.12801 |
| *C. russula* - ‘During’ | vs | *S. minutus* - ‘Before’ | 0.13091 | 0.13604 |
| *C. russula* - ‘After’ | vs | *S. minutus* - ‘During’ | 0.40765* | 0.45835* |
| *C. russula* - ‘After’ | vs | *S. minutus* - ‘Before’ | 0.38693* | 0.4209 |

The overlap of prey resources (measured using the Pianka’s index; Ojk) between *C. russula* and *S. minutus* is generally high at ∼37% to ∼50% overlap, depending on whether all MOTUs or core MOTUs are used (**Table 2**). These are significantly higher values than would be expected at random. *Crocidura russula* and *S. minutus* show a higher dietary overlap in Belle Île (All MOTUs Ojk = 0.4617, p < 0.05; Core MOTUs Ojk = 0.50289) than Ireland (All MOTUs Ojk = 0.35772, p < 0.01; Core MOTUs Ojk = 0.37765). When splitting the samples in Ireland according to invasion stage, *S. minutus* have a much higher resource overlap with *C. russula* ‘after’ the invasion (∼39–46%) compared to with *C. russula* ‘during’ the invasion (∼11–14%; **Table 2**). When accounting for all MOTUs, the level of overlap between *C. russula* ‘after’ the invasion and *S. minutus* is significantly higher than expected compared to simulated data (*S. minutus* ‘during’ Ojk = 0.40765, p = 0.0167; *S. minutus* ‘before’ Ojk = 0.38693, p = 0.028). This observation is consistent with the patterns shown in Figure 3.

## Discussion

The impact of the rapidly expanding invasive range of *C. russula* in Ireland demonstrates the need to consider both the spatial and temporal context of species invasions in a real-time setting. This study shows that an invasive species can quickly alter its behaviour and adapt to a new environment, leading to negative impacts that were not previously predicted based on co-existence of *C. russula* and *S. minutus* in other parts of their ranges. Here is an invasive species potentially exhausting local resources of larger prey taxa when it first enters an area and then beginning to shift its diet after becoming established towards the smaller prey taxa that are more essential for the survival of the native species. This interspecific competition then likely plays a key role in the subsequent disappearance of the native species in response to the invader.

In Ireland, the composition of the invasive *C. russula*’s diet changes between when it first invades an area and after it becomes established. Larger invertebrates such as worms (Haplotaxida), beetles (Coleoptera) and tough shelled millipedes (Glomerida) comprise a large portion of the *C. russula* diet at the invasion wavefront (‘during’), but is greatly reduced in the established ‘after’ invasion stage (**Fig. 2A**). The combination of high abundance, small territories and broad diet means that *C. russula* can exhaust local resources (Genoud, 1985). As a result, the level of interspecific dietary overlap increases from between 11–14% ‘during’ the invasion to between 39–46% ‘after’ the invasion (**Table 2**), and the NMDS plot shows a higher overlap between *S. minutus* generally and *C. russula* once the invader has become established (**Fig. 3**).

The invasion appears to be occurring in ‘layers’. Individuals from the first invading layer of *C. russula* (‘during’) are 20% larger in body mass (**Fig. 2B**) and longer (**Figs. S2 and S3**) than those ‘after’ the invasion. This may be the result of decreased intraspecific competition and/or investment in traits associated with increased dispersal abilities (Phillips *et al*., 2006; Burton *et al*., 2010). Their larger size may aid in their ability to predate on invertebrates that are too large for *S. minutus*, thus initially reducing competitive pressure. Previous studies have shown that shrews that differ greatly in size tend to have reduced niche overlap compared to shrew species closer in size (Churchfield and Rychlik, 2006). This would explain why both *C. russula* and *S. minutus* temporarily coexist at a restricted area where their ranges first meet (McDevitt *et al*., 2014). The relatively narrow niche width of *C. russula* (**Table S4**) provides some contradiction to previous claims that predation in shrews is likely opportunistic with little selection of prey (Castien and Gosalbez, 1999). The second established layer of *C. russula* (‘after’) are altering their diet to smaller prey taxa as a result of either exhausted resources, decrease in size (**Fig. 2B**) or both. Alternatively, the decrease in body mass in the second layer could suggest reduced energy intake from reduced food resources after the first layer (Seymour *et al*., 2005). This second layer of *C. russula* is what likely out-competes *S. minutus* for small prey resources that are key for their survival (Pernetta, 1976; Churchfield and Rychlik, 2006). This is why there is only a brief area of overlap, which eventually means they cannot coexist and *S. minutus* rapidly declines/disappears in as little as one year (McDevitt *et al*., 2014).

The dietary mechanisms by which this displacement occurs are subtle and reveal the importance of the DNA metabarcoding approach used here and its ability to identify prey tax beyond the order/family level (as is typical with morphology-based analysis of diet; Browett, O’Meara and McDevitt, 2020; Tournayre *et al*., 2020). The Pianka index identified considerable overlap in diet between these two shrews in both Ireland (up to 38%) and Belle Île (up to 50%; **Table 2**). This is supported by the NMDS plots (**Fig. 3**) and PERMANOVA showing minimal differences in the composition of prey between shrew species and country (**Table 1**). This level of dietary overlap has been seen between sympatric populations of water shrews *Neomys fodiens* and *S. minutus* in Poland (44% overlap), which was considered low for shrews (Churchfield and Rychlik, 2006). Overlap between the diet of sympatric shrews is considered high in general, and multi-species communities’ likely function as a result of subtle differences between habitat use and resource utilization (Churchfield and Sheftel, 1994). Therefore, the level of dietary overlap alone may be enough to explain their coexistence in Belle Île, but not Ireland. The POO values indicate that the majority of prey orders in Ireland are consumed by both predators (**Fig. 2A**). In contrast, there are key taxa that are consumed in Belle Île by one predator but not the other. There is an increased consumption of the orders Araneae, Hemiptera and Lepidoptera by Belle Île compared to Irish *S. minutus*, but not consumed by Belle Île *C. russula* (**Fig. 2** **and S8**). Instead, Belle Île *C. russula* have approx. 25% of their diet consisting of worms (Haplotaxida) and millipedes (Glomerida, Julida and Polydesmida), of which Belle Île *S. minutus* does not predate on. These prey orders may be key to providing competitive release between the shrews in Belle Île. Belle Île *S. minutus* also have a drastically increased consumption of Lepidoptera during the winter (**Fig. S10**), similar to previous observations of winter spikes of consuming Lepidopteran larvae using morphological approaches (Pernetta, 1976; Butterfield, Coulson and Wanless, 1981).

While DNA metabarcoding cannot identify life stage, it has identified a large proportion of this winter spike to be *Xestia xanthographa*. This moth species over-winters as nocturnal larvae (up to 35mm in size), feeding on various grasses (Skinner and Wilson, 2009). The nocturnal behaviour of *S. minutus* means they can take advantage of this slow moving and substantial food source during the less favourable winter conditions free from competition from *C. russula*. Another study in the Netherlands has also shown that partial niche segregation between *S. minutus* and the larger common shrew (*Sorex araneus*) over seasons may reduce interspecific competition (Ellenbroek, 1980). The small difference in prey taxa consumed by *S. minutus* in Belle Île between seasons suggests that they are predating on more readily available taxa between seasons, such as the apparent switch from Hemiptera in the summer to Lepidoptera in the winter (**Fig. S10**).

Another factor affecting resource use could be the morphology of the shrews themselves. Bite force and mechanical leverage of a shrew’s mandibles can determine the limits of prey size they can capture and consume (Cornette *et al*., 2015). Vega *et al*. (2016) examined the variation of shape and size of mandibles and skulls from *S. minutus* samples from various European regions including Ireland, Belle Île and multiple other islands. This study showed that *S. minutus* can exhibit morphological variability between different regions and islands in response to various environmental factors such as food availability and the presence of competitors. It also showed that the mandible size and shape of Irish *S. minutus* are distinct from other populations, likely a reflection of their long-term isolation from other European populations (Vega *et al.,* 2020). *Sorex minutus* from Belle Île are more similar to continental populations where they coexist with other species of shrews. The larger size of Belle Île individuals determined by this study (**Fig. 2B**) and the mandible shape inferred from Vega *et al*. (2016) may allow them to avail of a wider range of sizes of prey, which could explain the wider niche breadth measured by the Standardised Levin’s index (**Table S4**; Cornette *et al*., 2015). For example, species of Araneae consumed by Belle Île *S. minutus* are larger wolf spiders from the genera *Pardosa* and *Alopecosa* that can be up to 11mm in size, providing a substantial energy resource. Irish *S. minutus* show a reliance on smaller spiders such as *Pachygnatha* spp. measuring between 3–6mm (Nentwig *et al*., 2020).

## Conclusions

When the *C. russula* was first discovered in Ireland in 2007, it was thought that the species could potentially be a beneficial addition to the Irish mammalian fauna as a new prey resource for birds of prey, for example (Tosh *et al*., 2008). However, it quickly became apparent that the species was having a detrimental impact on the native shrew, *S. minutus* (Montgomery, Lundy and Reid, 2012; McDevitt *et al*., 2014; Montgomery, Montgomery and Reid, 2015). *Crocidura russula* is known to outcompete and displace other shrews on islands (Cornette *et al*., 2015) and here we have shown just how rapidly this can occur in real-time (McDevitt *et al*., 2014). Given that the eradication of an invasive shrew like *C. russula* on an island of Ireland’s size would not be logistically feasible (Seymour *et al*., 2005), this is obviously a concerning scenario for the island’s fauna. Future assessments of this shrew invasion should include different habitat types (e.g. woodland and peatland; Montgomery *et al*., 2015) to ascertain if coexistence with *S. minutus* is possible through differential habitat use as has been documented between other shrew species (Biedma *et al*., 2019; Keckel, Ansorge and Stefen, 2014). After being isolated for a long period of time in Ireland with the absence of larger competitors such as *S. araneus*, it has been suggested that *S. minutus* has altered their habitat use compared to their mainland counterparts (Michielsen, 1966) and it remains to be seen whether of the different habitat types in Ireland could support interspecific niche separation and act as potential refuges for *S. minutus* despite the presence of *C. russula* (McDevitt *et al*., 2014). However, Ireland has a relatively homogenous landscape and these potential refuge habitat types are sparse and doubts remain whether they could host a functioning metapopulation of *S. minutus* (McDevitt *et al*., 2014; Montgomery *et al*., 2015).

This obviously goes beyond the invasive shrew’s impacts on *S. minutus*. In terms of small mammal invasions on islands, there has been justifiably a lot of focus on the impacts caused by invasive commensals such as rats (*Rattus* spp.) and mice (*Mus* spp.) on other vertebrates (e.g. Jones *et al*., 2016), with perhaps less focus on their substantial impacts on invertebrates (St. Clair, 2011). In this study, we have shown that this invasive shrew initially preys on larger invertebrate taxa when they first invade an area before rapidly shifting towards smaller prey taxa after they become established. If indeed they are rapidly exhausting local resources of larger invertebrate species (Genoud, 1985), subsequent changes in terrestrial invertebrate communities can of course have severe impacts further downstream on ecosystem functioning and services (Sanchez-Bayo & Wyckhuys, 2019). It is therefore vital that further research is urgently undertaken (including pitfall trapping surveys (Oliver & Beattie, 1996) and improving local reference databases and primer optimisation for future DNA metabarcoding studies (Browett *et al.,* 2021; Curran *et al*., 2022)) to determine if *C. russula* is altering the composition of Ireland’s invertebrate community as its invasion rapidly progresses and what potential impacts this may have on the wider ecosystem on the island.

## Acknowledgements

Samuel SB was supported by a Pathway to Excellence PhD Scholarship from the University of Salford. Fieldwork was supported by grants from the Vincent Wildlife Trust and The Genetics Society awarded to Samuel SB and ADM. Laboratory work was supported by the NERC Environmental Omics Facility (grant reference NBAF1147) at the University of Sheffield awarded to ADM, Samuel SB and REA and a Pump Priming (University of Salford) award to ADM. We thank the members of the Molecular Ecology Group in Salford for advice on laboratory work and bioinformatics and Anthony Herrel for helping with the French licence. We would like to thank Gavin Horsburgh, Kathryn Maher and Rachel Tucker from the University of Sheffield for training and guidance during the laboratory work. Sequencing was performed by Tim Wright and the team at the Sheffield Diagnostic and Genetics Service, Sheffield Children’s Hospital NHS Foundation Trust. We thank Holly Broadhurst for providing illustrations of the shrews.

## Authors’ contributions

Allan D. McDevitt, Samuel S. Browett, Denise B. O’Meara, Jon M. Yearsley and Jeremy B. Searle conceived, and Samuel S. Browett, Allan D. McDevitt, Jon M. Yearsley, Rachael E. Antwis, Kevin J. Bown, Deborah A. Dawson and Owen S. Wangensteen designed the study. Samuel S. Browett, Stephen S. Browett and Rebecca Synnott performed the shrew sampling. Samuel S. Browett and Rebecca Synnott performed the laboratory work.

Samuel S. Browett analysed the data. Samuel S. Browett and Allan D. McDevitt wrote the manuscript and all authors contributed to editing, discussions and approval of the final manuscript.

## Data Accessibility

All bioinformatic steps and scripts can be found on github (https://github.com/ShrewlockHolmes/Browett_et_al_2022_BioRxiv). Raw sequence data will be made publicly available upon publication.

