## Supplementary Material for "Resource competition drives an invasion-replacement event among shrew species on an island"

### Methods

#### Trapping

Trap sites were chosen at hedgerows along secondary and tertiary roads adjacent to agricultural land (pasture or arable). Up to 100 Trip-traps (Proctor Bros. Ltd.) were placed, without bait, every 5m into long grass or mammal runs at the base of a single roadside hedgerow (**Figs. S1A and 1B**). One hedgerow may not produce enough samples, therefore each of the 9 zone/transect sample sites may consist of 2 or more sub-sample sites within 10km of each other. To avoid hunger related stress, traps were checked every 30 – 60 minutes. Non-target animals were released immediately.

#### DNA Extraction

Gut tracts were defrosted on ice, removed from ethanol and air dried. Gut contents were removed from the intestines and stomach using sterile instruments. Dissection was performed on disposable bench covers and tools were cleaned and flamed in between each sample to avoid cross-contamination. DNA was extracted from the gut contents using the Qiagen PowerSoil Kit, with protocol altered according to Alberdi *et al*. (2018). This kit removes PCR inhibitors that are associated with the digestive tract. Entire gut contents were weighed and added to the bead tubes as gut contents rarely amounted to the full recommended weight for the kit (0.3g). Some *C. russula* samples had more than 0.3g worth of gut contents. In such cases, the entire contents were manually homogenised in a 2ml Eppendorf, and the required amount was used for DNA extraction. Five extraction blanks were included.

DNA extractions were quantified using the Qubit broad range (BR) kit (Thermo Fisher Scientifc). DNA concentrations ranged from 43.9 ng/µl to 540 ng/µl. All DNA extractions were subsequently diluted in molecular grade water down to 10 – 15 ng/µl.

#### PCR

A 133bp fragment of the mtDNA COI gene was amplified from DNA extracts using the primers LepF1 (5′-ATTCHACDAAYCAYAARGAYATYGG-3′) and EPT-long-univR (5′-ACTATAAAARAAAATYTDAYAAADGCRTG-3′; Gillet *et al.*, 2015)) according to the protocol described in Browett *et al.* (2021). Gillet primers have issues with amplifying large amounts of host DNA (see results for details), but it was decided not to use vertebrate blocking primers because a) we want to detect if *C. russula* is predating on native vertebrates in Ireland and b) blocking primers could potentially block prey DNA to an extent (Vestheim, Deagle and Jarman, 2011). For multiplexing samples, a set of 24 unique eight base pair multiplex identifiers (MID) tags were added to the forward and reverse primers. This set of 24 primer pairs were arranged into 192 different combinations. Each sample was amplified in triplicate, with the five extraction blanks and 20 PCR blanks included. Samples were randomly distributed amongst 4 PCR plates, randomising host species, season, country, zone and transects to mitigate artificial inflation of inter-species/samples effects. Within each plate, 5 PCR blanks were included, randomly distributed amongst the plate, along with extraction blanks. Each one of these PCR plates constitutes a library of 77 to 86 samples, including negative controls. Each library was amplified in triplicate, but these PCR replicates were not individually barcoded (i.e. triplicates were pooled into a single representative sample). After PCR, triplicates were pooled and visually examined on a 1.2% agarose gel, stained with ethidium bromide. 2µl of sample were loaded into each well, with a 50bp ladder for reference. After amplification, each library was kept separate up to adapter ligation.

**Quantifying and Normalising PCR products**

To increase accuracy of normalising the full dataset, PCR products were quantified on a Fluorometer (FLUOstar OPTIMA), including between 6 and 7 standards. Numerous *S. minutus* samples showed amplification of regions larger than the 250bp Gillet region, causing complications directly comparing DNA concentrations with *C. russula* samples. Within each library, subsets of samples from the same species of shrews were normalised prior to DNA purification/bead clean.

**Bead Clean**

Before library preparation (i.e. the ligation of sequencing adapters onto PCR products), a bead clean was performed to purify the PCR products. To remove unwanted fragments smaller than 100bp, a left-side bead clean was performed using MAGBio HighPrep™ PCR Clean-up System beads at a 1.1 X ratio. To remove fragments larger than 300bp, a right-side bead clean was performed using a 0.8x ratio of beads to DNA template. Once the larger fragments have been removed from the *S. minutus* subsets (see Browett *et al.*,

2021), their concentrations are comparable to those of *C. russula* subsets. Each purified library subset was then quantified on the Fluorometer (FLUOstar OPTIMA), using 6 - 7 standards. Each subset was then pooled at equimolar concentration to form the 4 library pools. A second round of left side bead cleans were performed on each of the pools to remove any remaining primer dimer.

The success of each cleaning step was verified on an Agilent Tapestation using High Sensitivity screen tapes. Any fragments larger than the target amplicon (250bp) that remain at this point will not cause issues during the adapter ligation (library preparation) steps that follow. During library preparation, adapters will preferentially ligate to the smallest fragments available.

#### Adapter Ligation

Two sessions of adapter ligation were performed. The first session ligated adapters onto library 1 and 2, and the second session was to ligate adapters onto libraries 3 and 4. This was to keep each library with identical MID combinations separated from each other during library prep to mitigate tag-switching effects.

Adapters were ligated using the KAPA Hyper Prep Kit PCR-Free protocol with no modifications to the manufacturer's protocol. The NEXTFlex single index sequencing adapters for Illumina platforms were ligated onto each library. These adapters have a single 6bp index. A unique adapter index was associated with each of the 4 libraries, allowing the 328 samples to be multiplexed into a sequencing run.

To verify if adapters have successfully ligated and no un-ligated adapters remain, each library was examined on the TapeStation using the High Sensitivity screen tapes.

#### Sequencing

The libraries were quantified by qPCR using the KAPA library quantification kit for Illumina sequencing with 6 standards included. Each library was then diluted to 50nM and pooled at equal ratios to create a single equimolar pool at 50nM. This pool was then diluted to 8nM and clarified on another qPCR run using the same protocol. This pool was subsequently diluted to 4nM, containing all 303 samples, 5 extraction blanks and 20 PCR blanks. This library was checked again on the TapeStation using the High Sensitivity screen tapes before sequencing. The 4nM library was sequenced on two Illumina MiSeq runs using V2 300 cycle kits, both loaded aiming for 9pM with a 5% PhiX spike.

#### Bioinformatics

Processing of raw sequence reads was performed using the Obitools metabarcoding software pipeline (Boyer *et al*., 2016). After aligning the paired-end reads, sequences with an alignment quality score >40 and a length between 128–138bp were retained (Browett *et al*., 2021). All detected chimeras were removed using the uchime_denovo algorithm implemented in Vsearch (Rognes et al. 2016). Sequences were clustered using sumaclust (Boyer *et al.*, 2016) with a 98% similarity threshold (Alberdi *et al*., 2018; Browett *et al*., 2021) and singletons were removed. Sequence Molecular Operational Taxonomic Units (MOTUs) were taxonomically assigned using *blastn* against the NCBI database. Sequences required at least 80% identity and 90% alignment for a match (Frøslev *et al.*, 2017; Browett *et al.*, 2021). The top 25 matches were returned, and the most common taxid (taxonomy identifier) was assigned to that MOTU. MOTUs were required to have at least 98% identity for species-level assignment (Clare, Symondson & Fenton, 2014; Arrizabalaga-Escudero *et al*., 2018). MOTUs with high abundance and low taxonomic resolution were manually blasted against the Barcode of Life Database (BOLD Meiklejohn *et al.*, 2019) database to increase the resolution of the dataset. MOTUs between 97% and 98% were restricted to genus level assignments; between 96% and 97% were restricted to family level assignments and between 93% and 96% were restricted to order level assignments (Alberdi *et al.,* 2018). The maximum read count for each MOTU found in the 25 blanks was subtracted from the read counts of those MOTUs in each shrew sample (Elbrecht & Steinke, 2019).

### Results

#### Homogeneity of samples

When looking at samples grouped according to shrew species and country, the permutest showed a difference in dispersion/homogeneity between groups (permutest: $F = 8.831$, $p < 0.001$). Post-hoc pairwise permutest showed differences (pairwise permutest; p =

0.026) occurring with lower levels of dispersion in *C. russula* from Belle Île (mean dispersion = 0.61, SD = 0.12) and other groups (mean dispersion = 0.66, SD = 0.04), and lower dispersal in *C. russula* in Ireland (mean dispersion = 0.66, SD = 0.05) compared to Irish *S. minutus* (mean dispersion = 0.68, SD = 0.04; pairwise permutest; p = 0.042).

When grouping samples by species, country, season and zone, there was a difference between the homogeneity of group diet calculated using a permutest (F = 2.64, p = 0.004). Post-hoc pairwise permutest showed the differences occurring with lower levels of dispersal in *C. russula* groups (mean 0.62, SD = 0.09) compared to *S. minutus* groups (mean = 0.66, SD = 0.05; permutest p = 0.047). When grouping samples by species, country, season and transect, there was no significant difference between the dispersal calculated using the permutest method (F = 1.70, p = 0.061). This supports the sampling design that different transects may not influence differences between invasion zones in the PERMANOVA.

Comparison of quantification metrics

RRA, POO and wPOO performed very similarly for groups of samples (**Fig. S9**). However, there were discrepancies between the methods when quantifying the proportion of certain orders in the shrew diets. RRA returns higher proportion values for physically large orders of prey such as Stylommatophora (10mm – 70mm) in all populations and Coleoptera (5mm – 17mm) in Irish *S. minutus*. In the Belle Île *S. minutus* population, RRA returns high proportions of Araneae (2mm – 11mm) of which comprise species of larger bodied wolf spiders. Lepidoptera also returns high RRA values in Belle Île *S. minutus*, which could be a result of high predation of *Xestia xanthographa* larvae (up to 35mm) over the winter sampling period (see **Fig. S8**). Note that these orders are consumed by a large proportion of the populations (**Figs 2A and S8**).

### Diet Composition

Following recommendations by Deagle *et al*. (2019) to fully interpret the data, different methods were used and compared to determine the composition of the shrews' diets. The different metrics used here (RRA and POO) showed discrepancies between the importance of different food groups (**Fig. S9**). When using RRA to determine the composition of a consumer's diet, there are biological and technical biases to consider. Tissue from different prey species is digested by the consumer at different rates (Piñol, Senar and Symondson, 2019). Prey of smaller size have less DNA to be detected. RRA estimated lower levels of MOTUs classified under the Insecta class, which could be due to consumption of small species that results in fewer sequence reads. Some taxa groups and species are amplified better than others, resulting in a mismatch between proportions of input prey DNA before and after PCR amplification (Krehenwinkel *et al*., 2017; Bista *et al*., 2018). Feeding trials have shown a discrepancy of 3 times more or less the actual biomass reflected by relative read abundances (Thomas *et al*., 2016). Because of these biases, the diet reported at the population level will be using the POO metric here. The wPOO metric performed very similarly to POO and was thus not used. RRA was used in tandem to these analyses as a comparison.

Other than this study, only one other known study has used molecular techniques to determine the diet of *Crocidura* or *Sorex* shrews, which used only 5 samples of *S. minutus* in the UK (Ware *et al*., 2020). The majority of dietary analysis of *Crocidura* and *Sorex* species have relied on morphological analysis of gut contents (Pernetta, 1976; Churchfield and Sheftel, 1994; Churchfield and Rychlik, 2006; Brahmi *et al*., 2012) which are often restricted to classifying prey to order level. This is the largest molecular study of the diet of a *Sorex* and *Crocidura* species to date and offers new insights into the diets of these elusive mammals. DNA metabarcoding has shown that both shrews predate on a similar diversity of prey (**Table S4**), contradicting morphological methods that led to a consensus that *C. russula* predates on a wider variety of taxa than *S. minutus* (Churchfield, 2008). However, the larger size of *C. russula* means that while they can readily predate on smaller prey taxa such as springtails (Entomobryomorpha), a large proportion of the detectable diet here consists of relatively large prey groups such as worms (Haplotaxida) and tough shelled taxa such as snails (Stylommatophora) and millipedes (Julida, Glomerida, Polydesmida). There is huge variability between what individual shrews are eating, highlighting the importance of large sample sizes to characterise a shrew's diet (**Figs. S6 and S8**). A small number of reads originating from large livestock such as cattle were detected in the gut contents of multiple shrews. As the predation of cattle by shrews is of course highly unlikely, these reads likely occurred through secondary detection after shrews or prey invertebrates moved around mammalian dung. The low number of reads detected from other small mammals, such as *Clethrionomys glareolus*, do not definitively suggest predation of small mammals occurred. As a large number of *C. glareolus* were caught using the same traps during fieldwork, these reads are more likely a form of field contamination during trapping. This shows the primers ability to detect vertebrate DNA, but there was no clear evidence that *C. russula* has predated on any small vertebrates in Ireland or Belle Île as has been proposed in other regions (Brahmi *et al*., 2012).

Morphological identification methods have not typically suggested that *S. minutus* consume Stylommatophora (slugs and snails), likely due to their size (Pernetta, 1976; Churchfield and Rychlik, 2006). It should be acknowledged that soft bodied animals such as slugs may not be identifiable from morphological analysis (Deagle, Kirkwood and Jarman, 2009). DNA metabarcoding may provide the first evidence that slugs contribute to a significant portion of *S. minutus* diet. They are detected in over 50% of samples in both Ireland and Belle Île (**Fig. S8**) and have a POO of around 10% (**Fig. 2A**). The Stylommatophora species majorly contributing to *S. minutus* diet are relatively small (approx. 15 – 20mm long) and do not have any shells (e.g. *Deroceras laeve* and *Arion intermedius)* which would make predation easier for small shrews (**Fig. S7**). Secondary detection (i.e. detecting the food of the shrew's food) could possibly explain the detection of Stylommatophora if a shrew consumes an invertebrate that has come into contact with slug mucous or ingested slug tissue. The high number of reads coupled with the frequency of detection suggests that secondary detection is an unlikely reason for this result and *S. minutus* can actively predate on slugs. A small proportion of *S. minutus* samples in Ireland also contained reads from Haplotaxida (worms; **Figs 2A, S7 and S8**). This finding is similar to previous dietary assessments of *S. minutus* that concluded this to be opportunistic or scavenging behaviour due to the large size of worms and epigeal foraging behaviour of *S. minutus* (Churchfield and Rychlik, 2006).

### Supplementary Tables

**Table S1.** Sample size of shrews trapped in Ireland and Belle Île.

| *Season* | *Invasion Zone* | *Species* | *Sample Size* | | | |
|---|---|---|---|---|---|---|
| | | | West | South | East | **Total** |
| Summer | After | *C. russula* | 10 | 10 | 10 | **30** |
| | | *S. minutus* | - | - | - | **0** |
| | During | *C. russula* | 10 | 13 | 7 | **30** |
| | | *S. minutus* | 8 | 7 | 10 | **25** |
| | Before | *C. russula* | - | - | - | **0** |
| | | *S. minutus* | 10 | 10 | 10 | **30** |
| | Belle Île | *C. russula* | - | - | - | **20** |
| | | *S. minutus* | - | - | - | **20** |
| Winter | After | *C. russula* | 11 | 10 | 12 | **33** |
| | | *S. minutus* | - | - | - | **0** |

| | | | | | | |
|---|---|---|---|---|---|---|
| | During | *C. russula* | 11 | 10 | 10 | **31** |
| | | *S. minutus* | 9 | 8 | 5 | **22** |
| | Before | *C. russula* | - | - | - | **0** |
| | | *S. minutus* | 5 | 10 | 7 | **22** |
| | Belle Île | *C. russula* | - | - | - | **20** |
| | | *S. minutus* | - | - | - | **20** |
| Total | Ireland | *C. russula* | 42 | 43 | 39 | **124** |
| | | *S. minutus* | 32 | 35 | 32 | **99** |
| | Belle Île | *C. russula* | - | - | - | **40** |
| | | *S. minutus* | - | - | - | **40** |

**Table S2.** Values for the mean weights and total lengths of shrews trapped in various areas, with the standard deviations (SD) and standard errors (SE). N refers to the sample size.

| | | | | Weight (grams) | | | Length (mm) | | |
|---|---|---|---|---|---|---|---|---|---|
| **Species** | **Area** | **Season** | **N** | **Mean** | **SD** | **SE** | **Mean** | **SD** | **SE** |
| *C. russula* | Belle Ile | All | 41 | 9.841463 | 1.3015329 | 0.203265 | 111.8146 | 4.969636 | 0.776127 |
| *C. russula* | Ireland | All | 124 | 11.008065 | 1.7227128 | 0.154704 | 115.8432 | 4.376277 | 0.393001 |
| *C. russula* | After | All | 63 | 10.57619 | 1.3668559 | 0.172208 | 114.0119 | 3.502104 | 0.441224 |
| *C. russula* | During | All | 61 | 11.454098 | 1.9378316 | 0.248114 | 117.7344 | 4.407943 | 0.564379 |
| *S. minutus* | Belle Ile | All | 41 | 4.360976 | 0.7844992 | 0.122518 | 96.47073 | 5.353468 | 0.836071 |
| *S. minutus* | Ireland | All | 100 | 3.344 | 0.4511086 | 0.045111 | 91.417 | 4.539814 | 0.453981 |
| *S. minutus* | Before | All | 52 | 3.284615 | 0.4202635 | 0.05828 | 90.62115 | 4.431852 | 0.614587 |
| *S. minutus* | During | All | 48 | 3.408333 | 0.4783986 | 0.069051 | 92.27917 | 4.543078 | 0.655737 |
| *C. russula* | Belle Ile | Summer | 20 | 8.83 | 0.7773775 | 0.173827 | 109.33 | 4.408616 | 0.985797 |

| | | | | | | | | | |
|---|---|---|---|---|---|---|---|---|---|
| *C. russula* | Belle Ile | Winter | 21 | 10.804762 | 0.9035907 | 0.19718 | 114.181 | 4.344838 | 0.948121 |
| *C. russula* | During | Summer | 30 | 10.766667 | 1.6589015 | 0.302873 | 116.0067 | 4.326895 | 0.789979 |
| *C. russula* | During | Winter | 31 | 12.119355 | 1.9799696 | 0.355613 | 119.4065 | 3.858492 | 0.693006 |
| *C. russula* | After | Summer | 30 | 10.506667 | 1.2687037 | 0.231633 | 114.3717 | 3.79518 | 0.692902 |
| *C. russula* | After | Winter | 33 | 10.639394 | 1.4671272 | 0.255394 | 113.6849 | 3.237005 | 0.56349 |
| *S. minutus* | Belle Ile | Summer | 20 | 3.71 | 0.2693071 | 0.060219 | 92.28 | 3.486757 | 0.779663 |
| *S. minutus* | Belle Ile | Winter | 21 | 4.980952 | 0.5784627 | 0.126231 | 100.4619 | 3.388433 | 0.739417 |
| *S. minutus* | Before | Summer | 30 | 3.27 | 0.1784029 | 0.032572 | 88.31333 | 3.559033 | 0.649788 |
| *S. minutus* | Before | Winter | 22 | 3.304545 | 0.6198904 | 0.132161 | 93.76818 | 3.496309 | 0.745416 |
| *S. minutus* | During | Summer | 25 | 3.384 | 0.2115026 | 0.042301 | 89.024 | 2.52888 | 0.505776 |
| *S. minutus* | During | Winter | 23 | 3.434783 | 0.6623709 | 0.138114 | 95.81739 | 3.46235 | 0.72195 |

**Table S3**. Breakdown of number of sequences retained through quality filtering steps. Sample size does not include controls. Last column is blank for individual libraries as libraries were combined prior to singleton removal.

| *Library* | *Sample Size* | *Sequencing Run* | *Reads Generated* | *Quality Filter* | *Correct Length* | *Chimera removal* | *Singleton Removal* |
|---|---|---|---|---|---|---|---|
| 1 | 78 | 1 | 3961614 | 3104266 | 2883535 | 2883535 | NA |
| 1 | 78 | 2 | 4221401 | 3300256 | 3065428 | 3065428 | NA |
| 2 | 80 | 1 | 3589389 | 2679527 | 2418157 | 2418157 | NA |
| 2 | 80 | 2 | 3918315 | 2911989 | 2629218 | 2629218 | NA |
| 3 | 70 | 1 | 2956215 | 2223848 | 2042631 | 2042631 | NA |
| 3 | 70 | 2 | 3188761 | 2407114 | 2212002 | 2212002 | NA |
| 4 | 77 | 1 | 4029211 | 3135357 | 2826117 | 2826117 | NA |
| 4 | 77 | 2 | 4307512 | 3340485 | 3014415 | 3014415 | NA |
| Total | 305 | NA | 30172418 | 23102842 | 21091503 | 21091503 | 21045832 |

**Table S4**. Alpha diversity values for groups of shrews according to species and country of origin. All groups were randomly subsampled to the smallest sample size (N = 23) 50 times and the average value was taken to make each group more comparable.

| Group | Prey Taxa Level | Sample Size | Mean Richness | Richness sd | Mean Shannon Index | Shannon Index sd | Mean Levin's Index | Levin's Index sd | Mean Standardised Levin's Index | Standardised Levin's sd |
|---|---|---|---|---|---|---|---|---|---|---|
| *C. russula* - Belle Ile | MOTU | 23 | 213 | 0.00 | 5.03 | 0.00 | 101.71 | 0.00 | 0.48 | 0.00 |
| *C. russula* - Ireland | MOTU | 23 | 196.52 | 13.85 | 5.05 | 0.08 | 115.42 | 10.76 | 0.58 | 0.03 |
| *S. minutus* - Belle Ile | MOTU | 23 | 181.16 | 7.51 | 4.97 | 0.05 | 105.73 | 6.20 | 0.58 | 0.02 |
| *S. minutus* - Ireland | MOTU | 23 | 216.06 | 14.96 | 5.10 | 0.09 | 108.72 | 13.99 | 0.50 | 0.05 |
| *C. russula* - Belle Ile | species | 23 | 129 | 0.00 | 4.44 | 0.00 | 56.86 | 0.00 | 0.44 | 0.00 |
| *C. russula* - Ireland | species | 23 | 116.58 | 6.92 | 4.46 | 0.07 | 64.72 | 4.89 | 0.55 | 0.03 |
| *S. minutus* - Belle Ile | species | 23 | 138.12 | 3.91 | 4.64 | 0.04 | 74.63 | 3.51 | 0.54 | 0.02 |
| *S. minutus* - Ireland | species | 23 | 151.76 | 11.01 | 4.67 | 0.08 | 68.92 | 6.24 | 0.45 | 0.03 |
| *C. russula* - Belle Ile | genus | 23 | 104 | 0.00 | 4.19 | 0.00 | 44.72 | 0.00 | 0.42 | 0.00 |

| | | | | | | | | | | |
|---|---|---|---|---|---|---|---|---|---|---|
| *C. russula* - Ireland | genus | 23 | 92.02 | 5.24 | 4.15 | 0.06 | 43.80 | 3.13 | 0.47 | 0.02 |
| *S. minutus* - Belle Ile | genus | 23 | 119.42 | 4.21 | 4.49 | 0.04 | 65.52 | 3.42 | 0.54 | 0.02 |
| *S. minutus* - Ireland | genus | 23 | 124.6 | 7.40 | 4.41 | 0.07 | 54.03 | 4.63 | 0.43 | 0.03 |
| *C. russula* - Belle Ile | family | 23 | 78 | 0.00 | 3.92 | 0.00 | 34.98 | 0.00 | 0.44 | 0.00 |
| *C. russula* - Ireland | family | 23 | 67.8 | 3.63 | 3.79 | 0.06 | 29.83 | 2.08 | 0.43 | 0.02 |
| *S. minutus* - Belle Ile | family | 23 | 79.74 | 1.82 | 4.09 | 0.03 | 47.58 | 1.67 | 0.59 | 0.02 |
| *S. minutus* - Ireland | family | 23 | 80.32 | 4.42 | 3.98 | 0.06 | 38.86 | 2.83 | 0.48 | 0.03 |
| *C. russula* - Belle Ile | order | 23 | 30 | 0.00 | 3.10 | 0.00 | 18.52 | 0.00 | 0.60 | 0.00 |
| *C. russula* - Ireland | order | 23 | 26.22 | 1.53 | 2.90 | 0.04 | 14.27 | 0.65 | 0.53 | 0.04 |
| *S. minutus* - Belle Ile | order | 23 | 26.16 | 1.09 | 2.91 | 0.03 | 15.45 | 0.47 | 0.57 | 0.02 |
| *S. minutus* - Ireland | order | 23 | 24.66 | 1.61 | 2.87 | 0.05 | 15.06 | 0.68 | 0.60 | 0.04 |

### Supplementary Figures

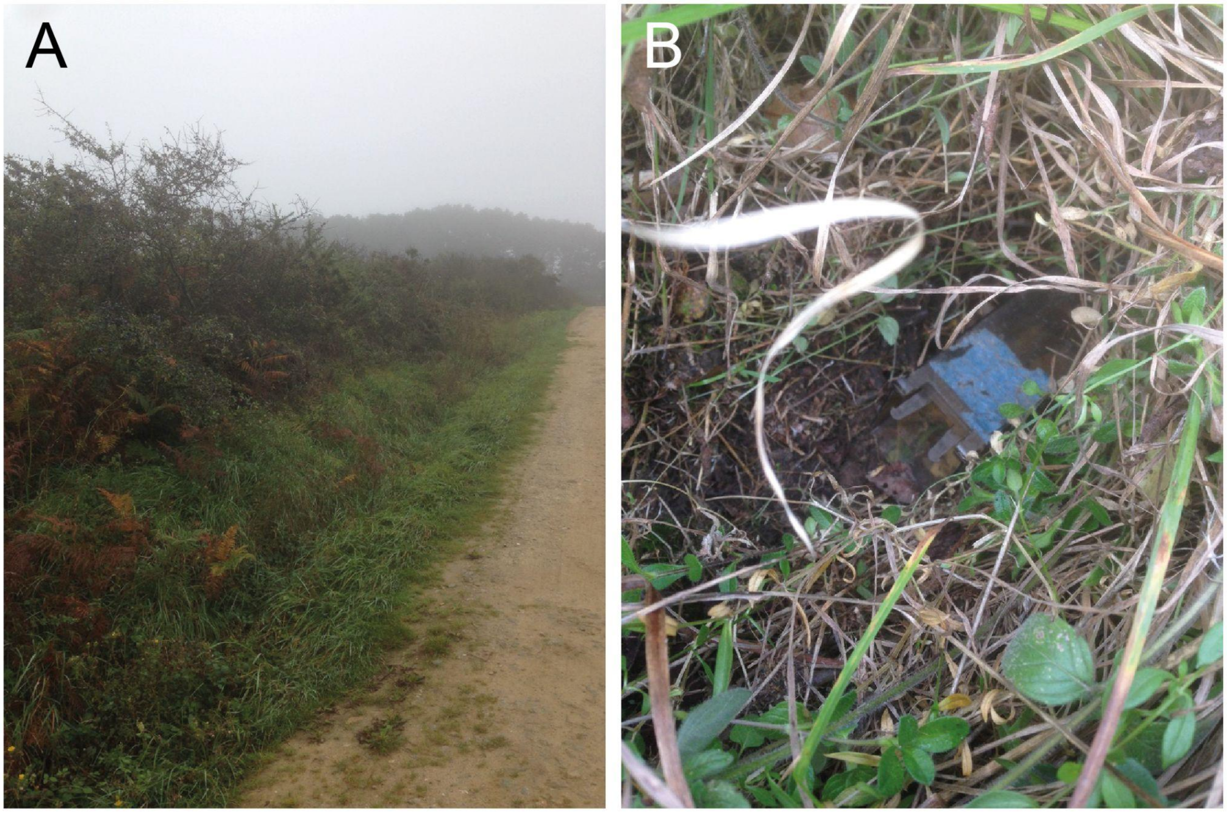


**Figure S1.** Typical areas for placing Trip-traps in hedgerows along roads (A) and a close-up of Trip-trap placement.

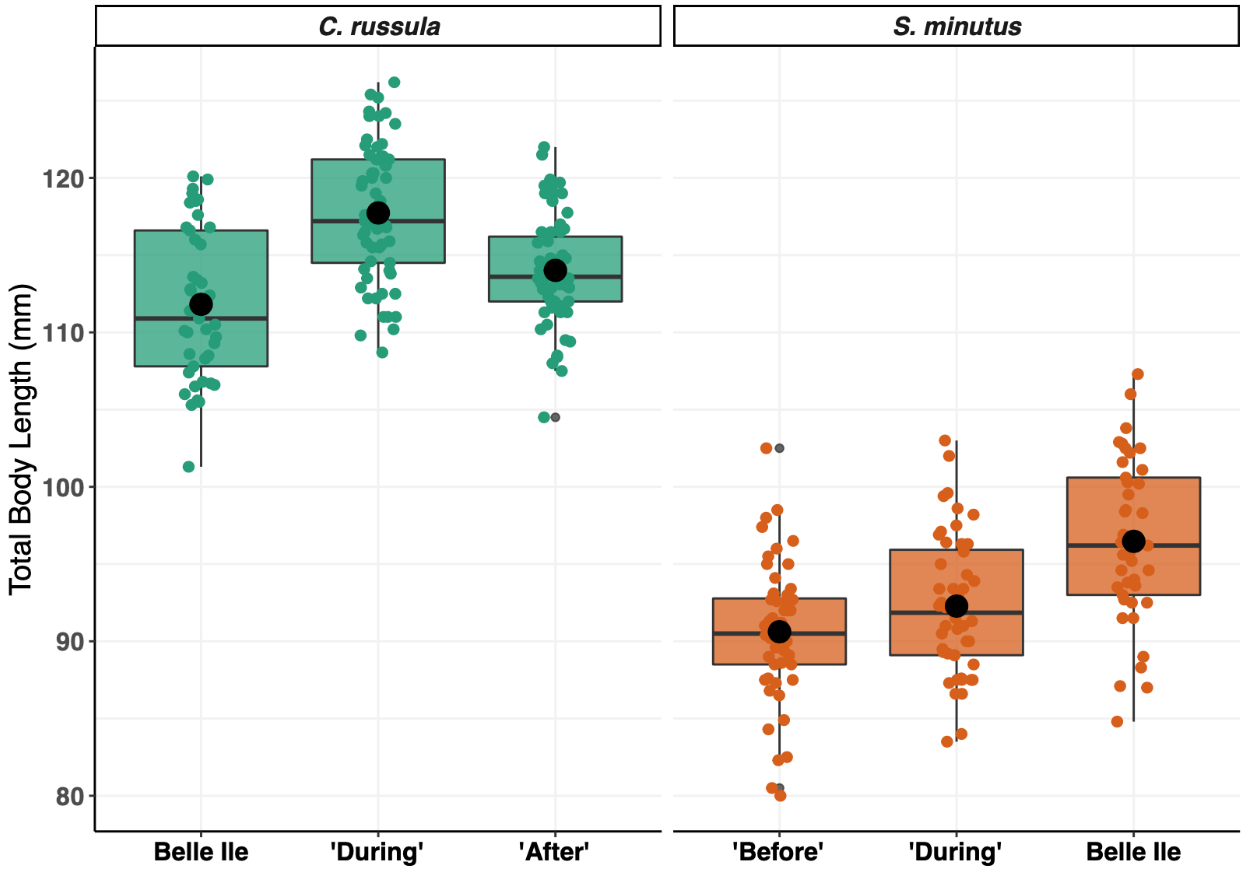


**Figure S2.** The total body lengths of *C. russula* (green) and *S. minutus* (orange) for each invasion stage in Ireland (‘Before’, ‘During’ and ‘After’; see main text) and the ‘Control’ site in Belle Île. The black dots represent the mean value for each group.

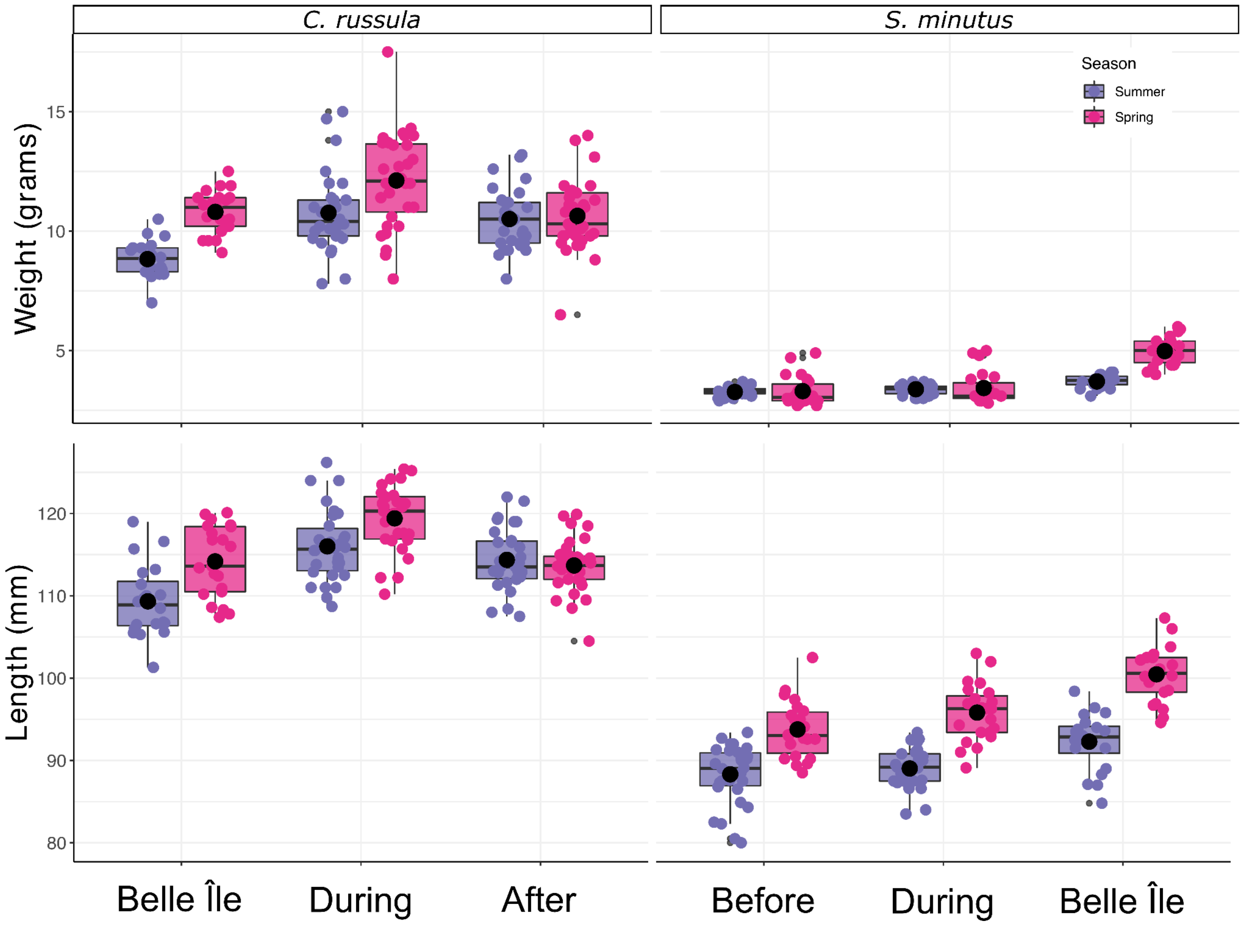


**Figure S3.** The weight (grams) and total body lengths (mm) of *C. russula* (left plots) and *S. minutus* (right plots) for each invasion stage in Ireland ('Before', 'During' and 'After') and the 'Control' site in Belle Île by season (summer in purple and spring in pink). The black dots represent the mean value for each group.

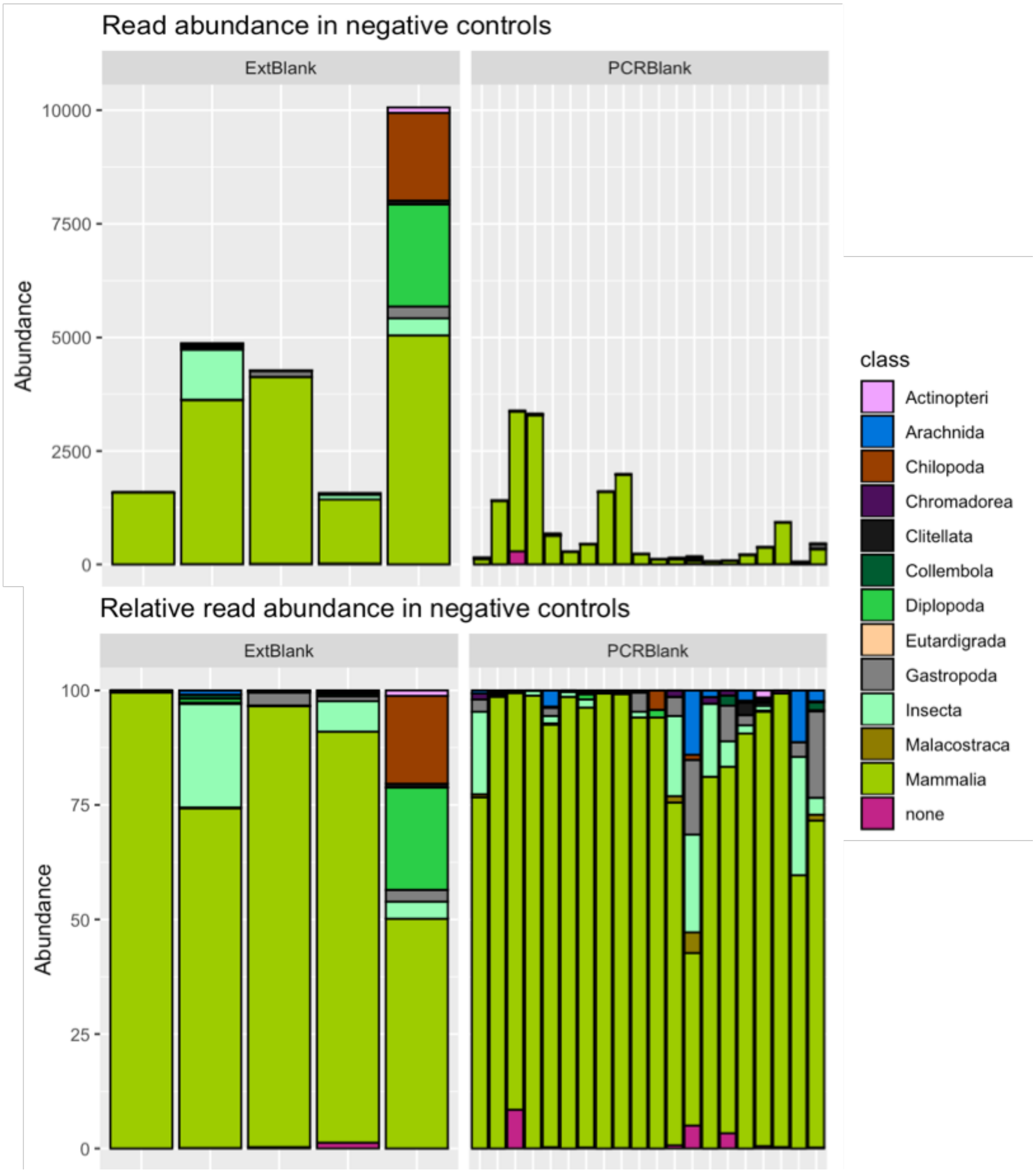


**Figure S4.** Read numbers in the negative controls. The top panel is total read count, while the bottom is the relative read abundance. Each vertical bar represents an individual negative control sample. Colour codes correspond to taxonomic assignment and 'none' represents reads that could not be taxonomically assigned.

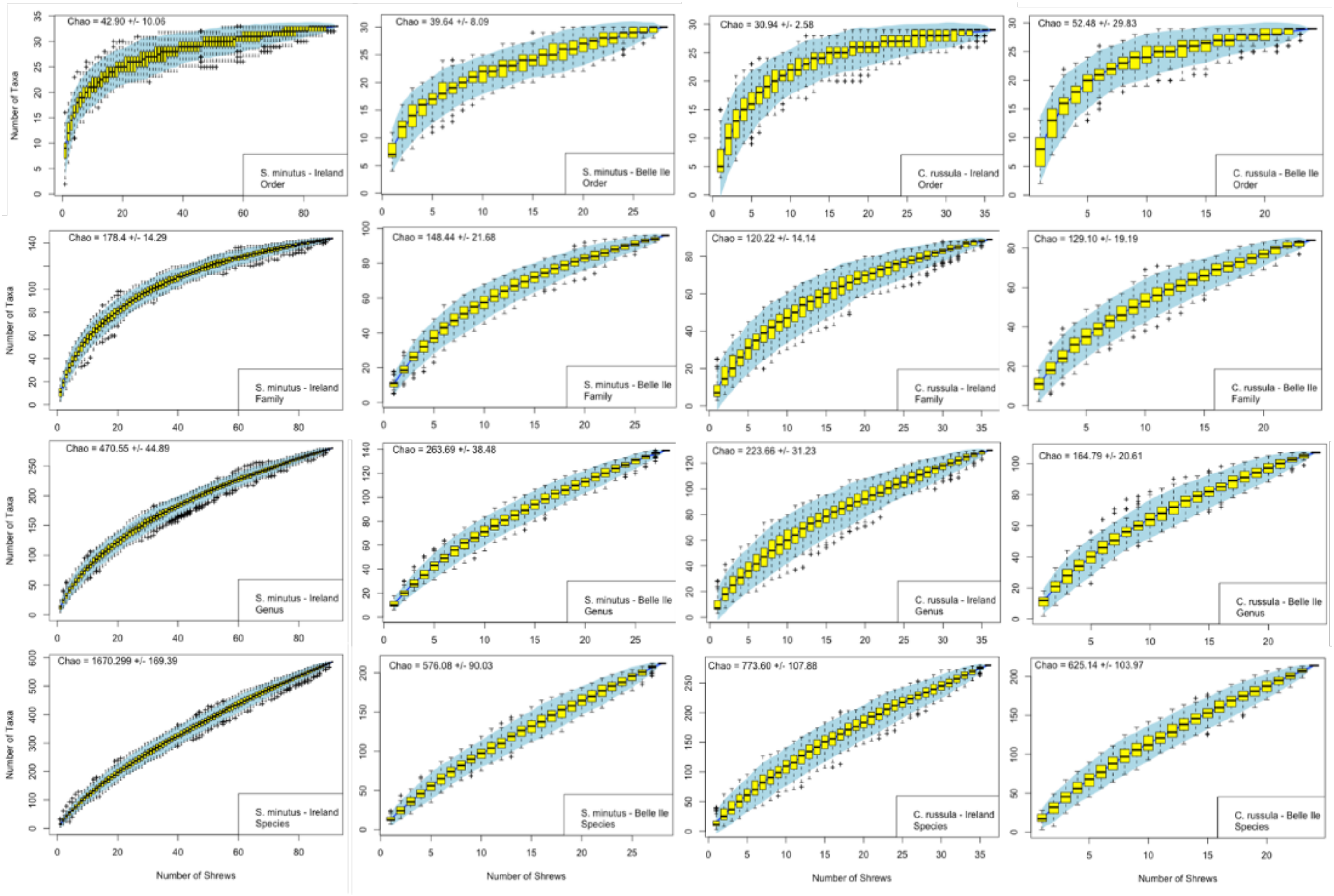


**Figure S5**. Species accumulation curves for (from left to right) *S. minutus* in Ireland, Belle Île and *C. russula* from Ireland and Belle Île. From bottom to the top, taxa have been kept at MOTU/species level and agglomerated to genus, family and order level.

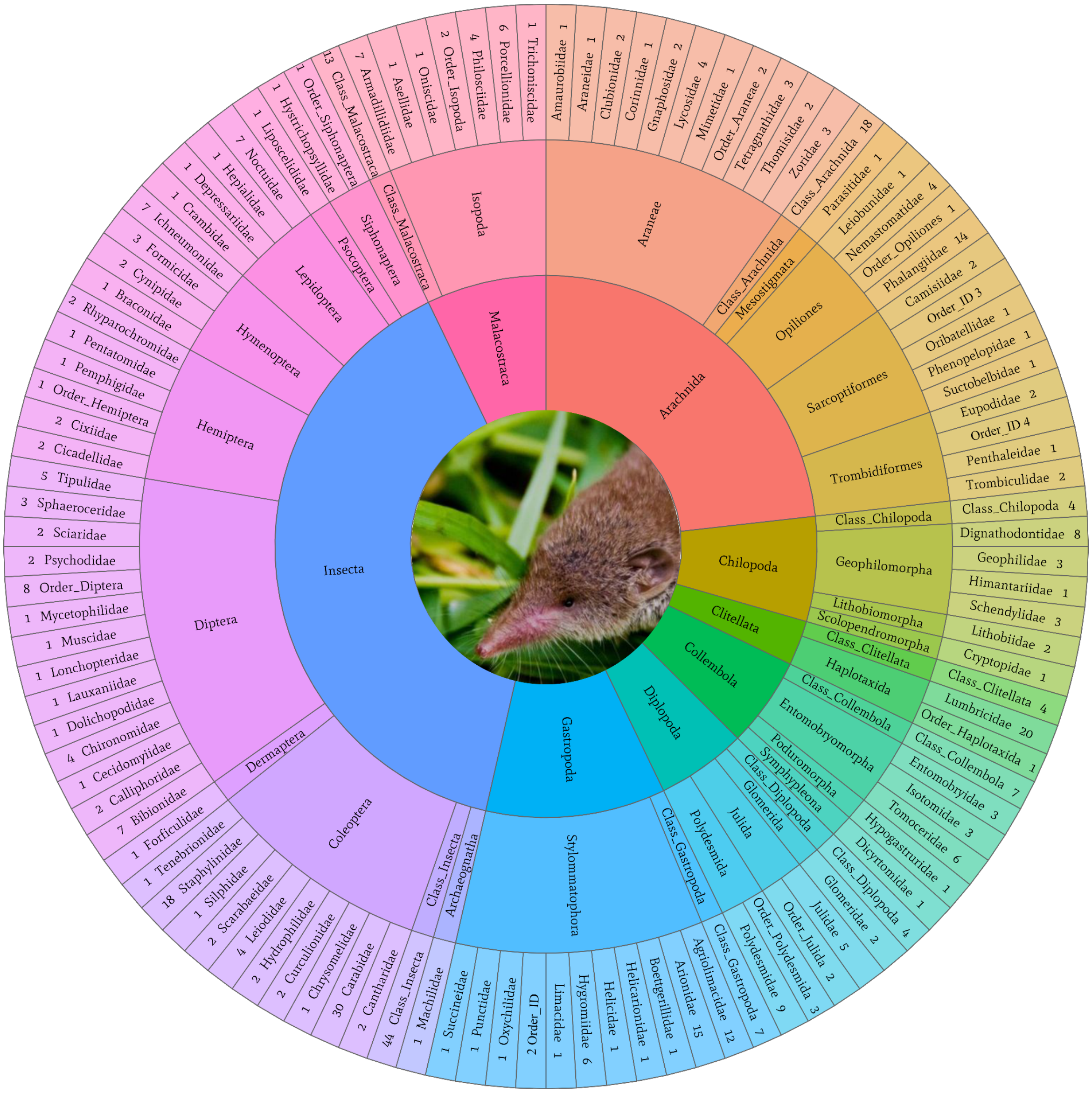


**Figure S6.** Range of prey taxa identified in *Crocidura russula*. A total of 438 Molecular Operational Taxonomic Units (MOTUs) were detected in *C. russula*. The layers of the taxonomic wheel move from inside to the outside showing class, order and family level. The numbers represent the number of MOTUs found in each group. 'Class ID' and 'Order ID' indicate the number of MOTUs only identifiable to class and order level, respectively.

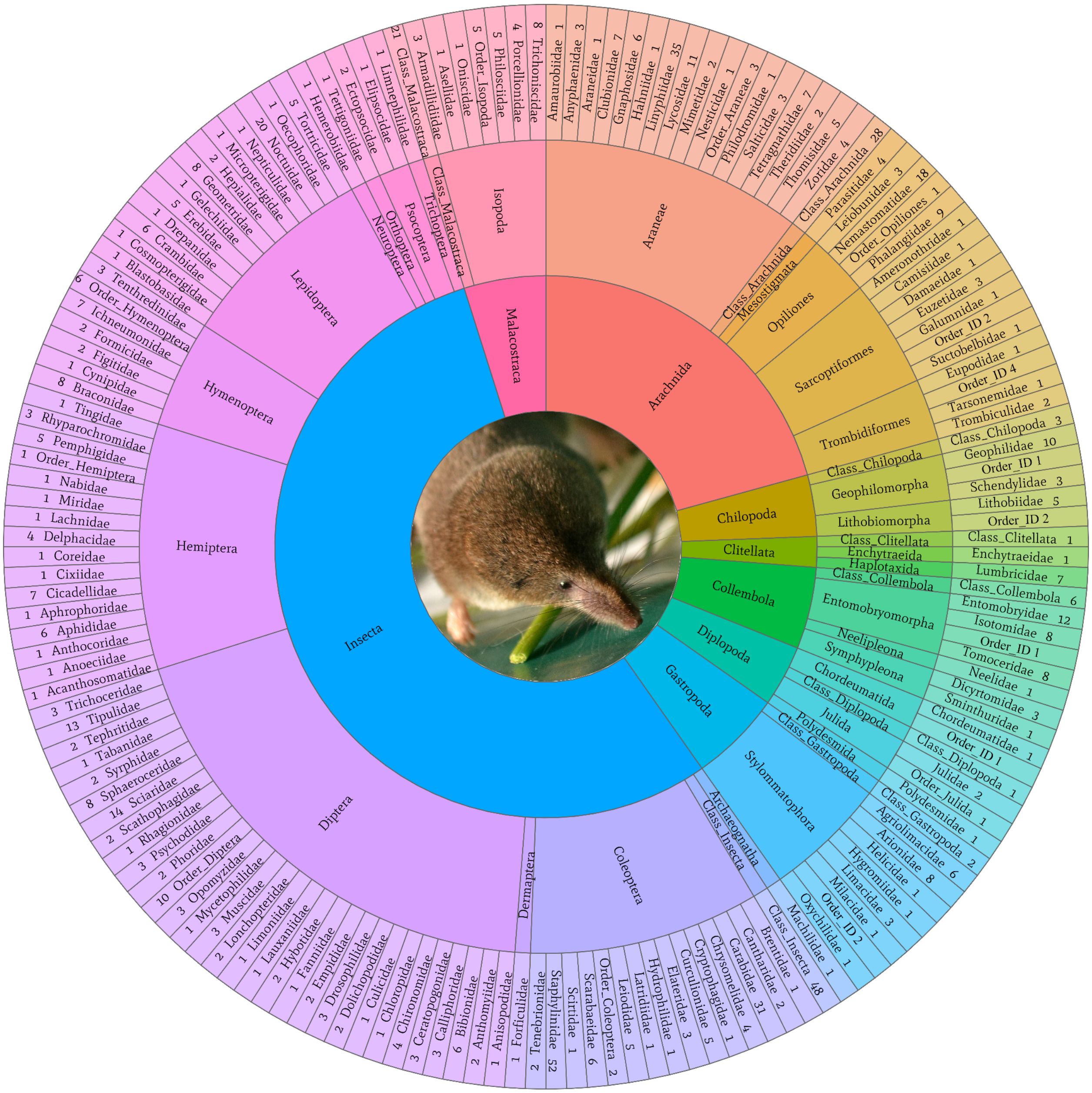


**Figure S7.** Range of prey taxa identified in *Sorex minutus*. A total of 721 Molecular Operational Taxonomic Units (MOTUs) were detected in *S. minutus*. The layers of the taxonomic wheel move from inside to the outside showing class, order and family level. The numbers represent the number of MOTUs found in each group. ‘Class ID’ and ‘Order ID’ indicate the number of MOTUs only identifiable to class and order level, respectively.

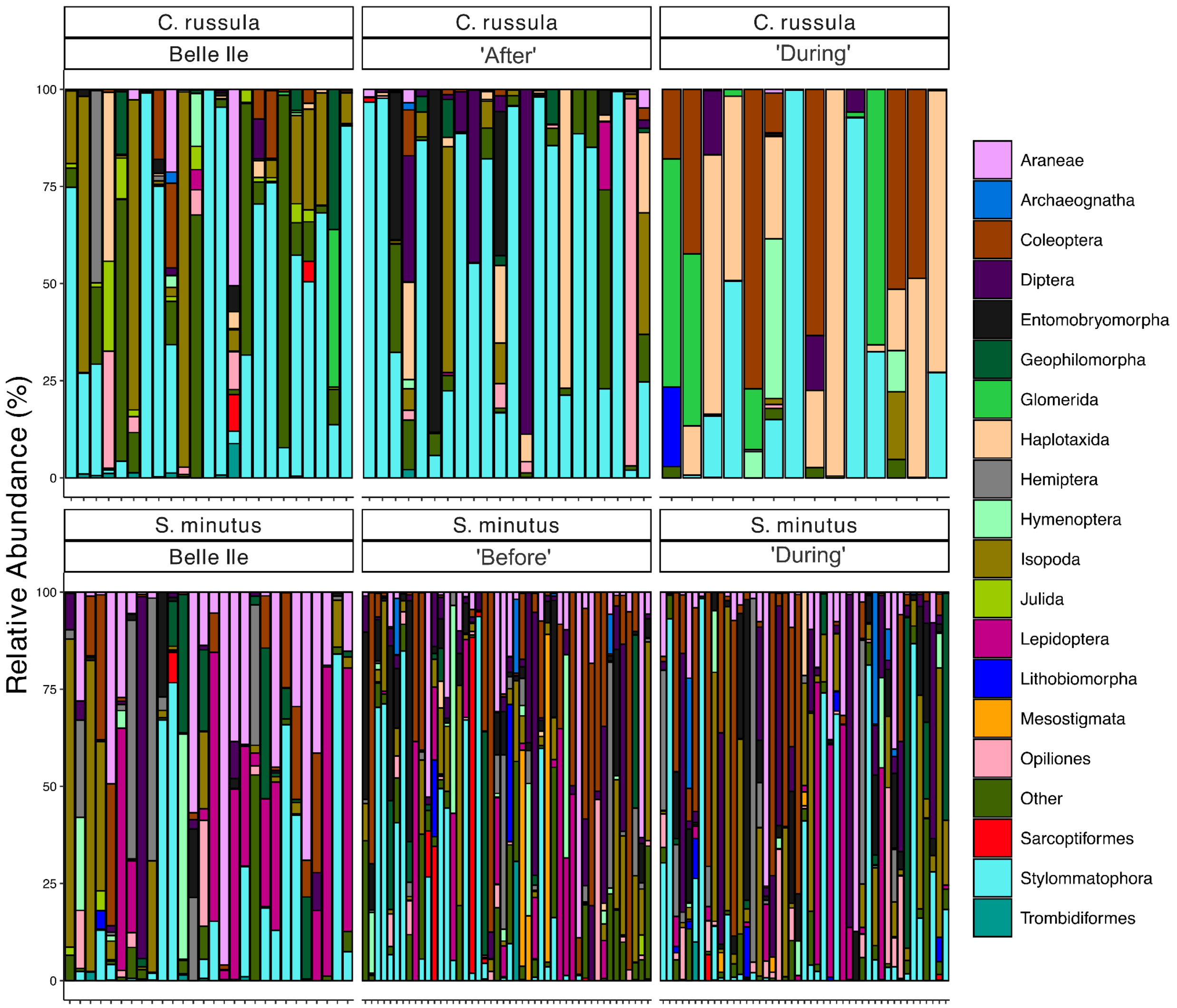


**Figure S8.** The Relative Read Abundance (RRA) of prey order in each sample, sectioned according to species, country and invasion stage ('Before, 'During' and 'After'; see main text). Each vertical bar represents an individual shrew. The colour represents the proportion of the reads found for that shrew that correspond to the prey order of the same colour.

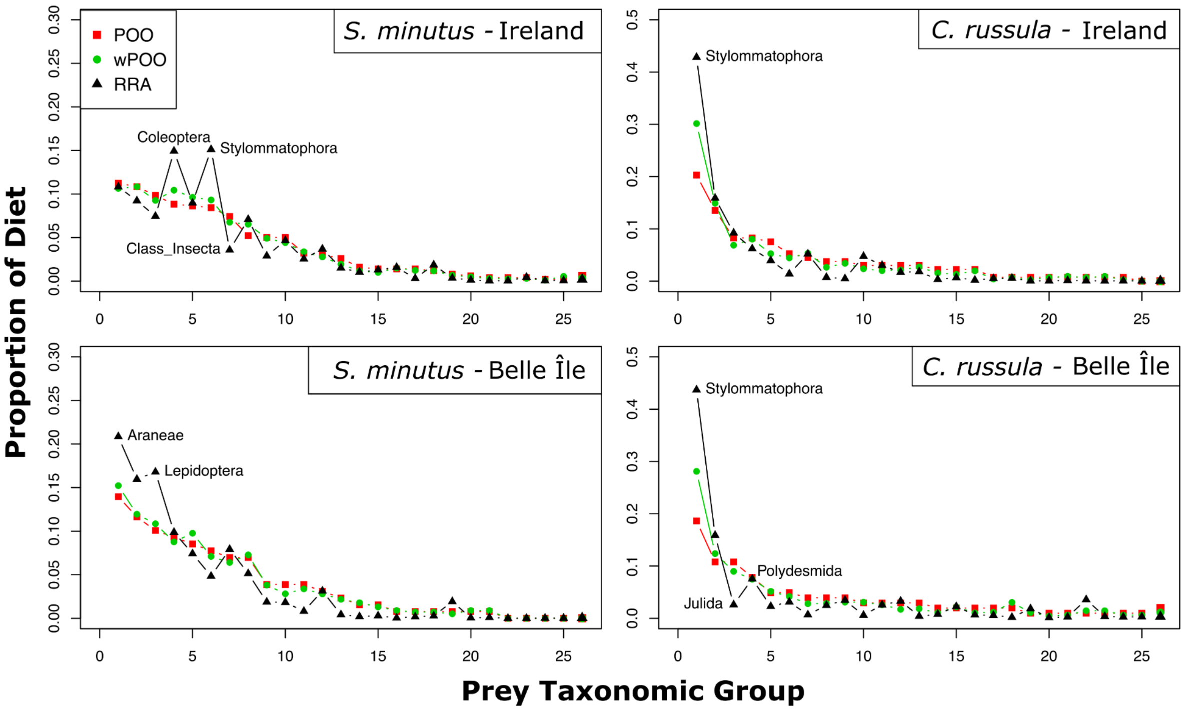


**Figure S9.** Comparison of methods to estimate proportion of diets from each taxa group (at order level). Certain groups where relative read abundance (RRA) disagrees with Percentage of Occurrence (POO) and wPOO estimates are labelled. For example, RRA (black) potentially overestimates the proportion of the diet consisting of Stylommatophora compared to POO (red) and wPOO (green).

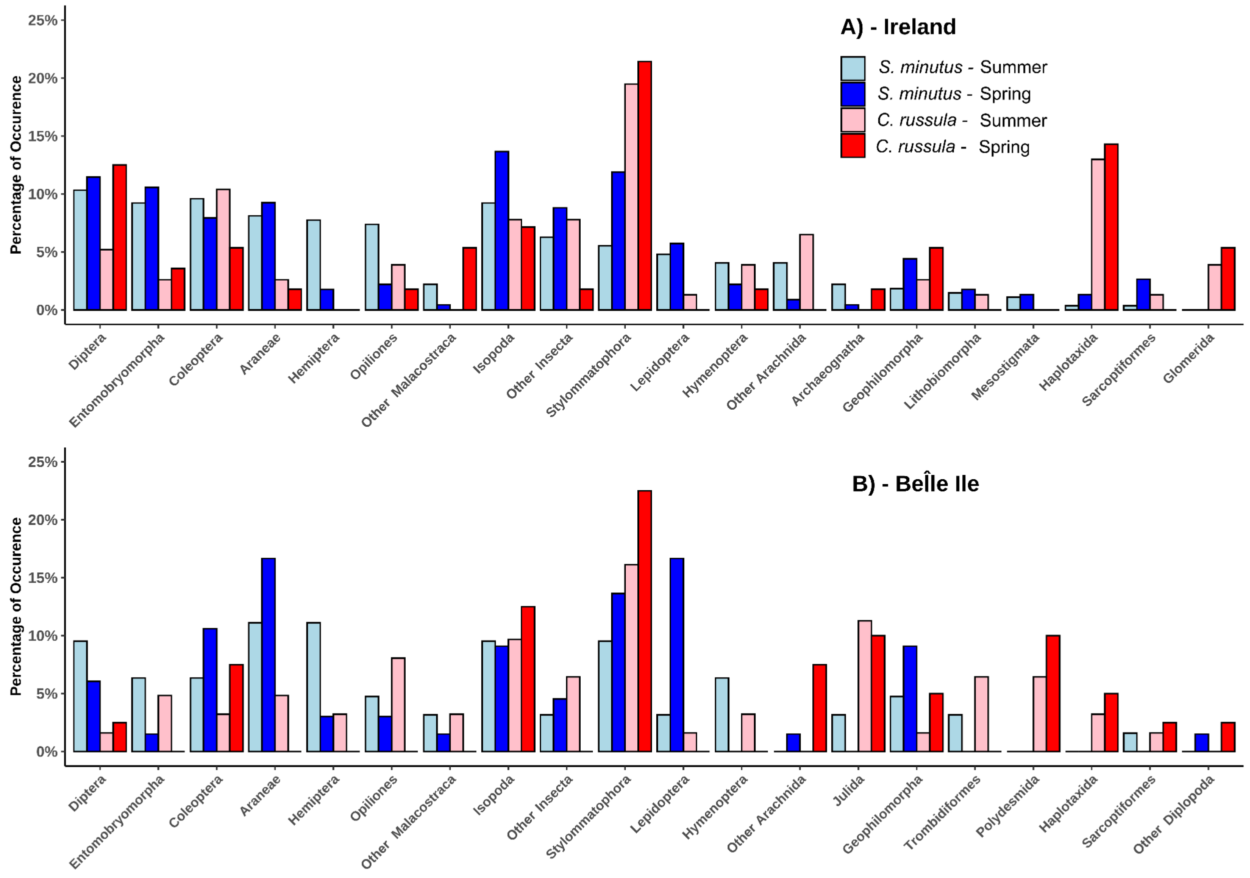


**Figure S10.** Percentage of Occurrence (POO) of top 20 orders (some only identified to class level). This acts to show the proportion of the diet each order contributes to. Groups are split into species and season in Ireland (top) and Belle Île (bottom).
